# Engineering protease-resistant peptides to inhibit human parainfluenza viral respiratory infection

**DOI:** 10.1101/2021.02.22.432312

**Authors:** Victor K. Outlaw, Ross W. Cheloha, Eric M. Jurgens, Francesca T. Bovier, Yun Zhu, Dale F. Kreitler, Olivia Harder, Stefan Niewiesk, Matteo Porotto, Samuel H. Gellman, Anne Moscona

## Abstract

The lower respiratory tract infections affecting children worldwide are in large part caused by the parainfluenza viruses (HPIVs), particularly HPIV3, along with human metapneumovirus and respiratory syncytial virus, enveloped negative-strand RNA viruses. There are no vaccines for these important human pathogens, and existing treatments have limited or no efficacy. Infection by HPIV is initiated by viral glycoprotein-mediated fusion between viral and host cell membranes. A viral fusion protein (F), once activated in proximity to a target cell, undergoes a series of conformational changes that first extend the trimer subunits to allow insertion of the hydrophobic domains into the target cell membrane, and then refold the trimer into a stable postfusion state, driving the merger of the viral and host cell membranes. Lipopeptides derived from the C-terminal heptad repeat (HRC) domain of HPIV3 F inhibit infection by interfering with the structural transitions of the trimeric F assembly. Clinical application of this strategy, however, requires improving the *in vivo* stability of antiviral peptides. We show that the HRC peptide backbone can be modified via partial replacement of α-amino acid residues with β-amino acid residues to generate α/β-peptides that retain antiviral activity but are poor protease substrates. Relative to a conventional α-lipopeptide, our best α/β-lipopeptide exhibits improved persistence *in vivo* and improved anti-HPIV3 antiviral activity in animals.

## Introduction

Human parainfluenza viruses (HPIV1-4) are paramyxoviruses that cause human respiratory diseases, including 30-40% of childhood croup, bronchitis, bronchiolitis, and pneumonia. Infants, children, and immune-compromised individuals are particularly vulnerable to HPIV infection^1,2^, and no FDA-approved vaccines or targeted antiviral therapeutics are currently available for any of the HPIV serotypes. Vaccines for parainfluenza have long been under study, and a limited number of vaccines are in Phase I clinical trials^3,4^; however, no vaccine is close to FDA approval, and even this eventual development will not eliminate HPIV3 disease in the foreseeable future. While corticosteroids have decreased hospitalizations for HPIV1-associated croup^5^, HPIV2 and HPIV3 infections remain untreatable, and HPIV3 is responsible for more hospitalizations than HPIV1 and 2 combined.^6^

As for other paramyxoviruses, HPIV3 infection is initiated by the fusion of viral and host cell membranes, mediated by attachment (HN for HPIV3; H or G for related paramyxoviruses) and fusion (F) glycoproteins, that form the fusion complex required for entry.^7-17^ HPIV3 HN attaches to the host cell by engaging a surface receptor. The homotrimeric HPIV3 F, initially anchored to the viral envelope by transmembrane segments and adjacent C-terminal heptad repeat (HRC) domains in a metastable prefusion conformation, is then activated by the receptor-engaged HN and undergoes a sequence of structural transitions. First, N-terminal heptad repeat (HRN) domains within F extend, and fusion peptide segments insert into the host cell membrane to form a transient prehairpin intermediate. Subsequent refolding of F causes the HRN and HRC domains to form a highly stable six-helix bundle (6HB), a process that is coupled to membrane fusion.

HRC peptides can inhibit fusion by binding to the transiently exposed HRN segments of the short-lived prehairpin intermediate and preventing 6HB formation.^18,19^ We have previously described fusion inhibitory peptides derived from the HPIV3 F HRC domain that display potent antiviral activity against HPIV3, as well as against other paramyxoviruses.^20-30^ Conjugating cholesterol to an HRC peptide enhances antiviral potency by targeting the lipopeptide to the plasma membrane, the site of fusion.^23-25,31^ The design strategy discussed here is aimed at enhancing antiviral activity of HPIV3-derived HRC peptides *in vivo* by fostering resistance to degradation by proteases.

Rapid degradation by proteases and renal clearance hinder the development of peptide-based drugs.^32^ Our approach features two complementary strategies, lipid conjugation and β-amino acid incorporation, to circumvent these problems. We have shown that attaching a lipophilic unit to peptides comprised entirely of α-amino acid residues (α-peptides) improves pharmacokinetic properties of fusion-inhibitory peptides, decreases protease sensitivity, and increases antiviral potency by several logs relative to unmodified α-peptides.^23,25,26^ In separate studies, we have demonstrated that replacing a subset of α-amino acid residues with β-amino acid residues can generate an “α/β-peptide” with substantially reduced susceptibility to proteolysis relative to an analogous α-peptide.^33-35^ These α/β-peptides can structurally and functionally mimic α-helical α-peptides from which they were derived.^36-38^ Here we show that combining lipid conjugation and α-to-β substitution results in decreased susceptibility to proteolytic degradation, enhanced pharmacokinetic properties, and potent anti-HPIV3 activity *in vivo*. This study provides the first evidence that backbone-modified fusion-inhibitory peptides can display activity in animals.

## Results

### Design of α/β-peptide analogues of α-VI

Our starting point was a conventional peptide based on the HPIV3 F HRC domain that contains two substitutions, E459V^21,23^ and A463I^23^, designated “α-VI” (Figure 1A). A series of α/β-peptide derivatives was examined (Figure 1A). Our design strategy was based on the heptad sequence repeat pattern inherent in HRC-derived peptides. One heptad, the positions of which are designated *a-g*, corresponds approximately to two turns of an α-helix. A “helix wheel” diagram depicts the positions of the seven side chains projecting from a single heptad based on a view along the helix axis (Figure 1B). HRC domain side chains at positions *a, d, e* and *g* typically contact the core HRN trimer, while side chains at *b, c* and *f* generally lack interactions with other components of the 6HB. Our design strategy placed β residues at non-contact positions in order to minimize disruption of the HRC-HRN interface.

**Figure 1.**
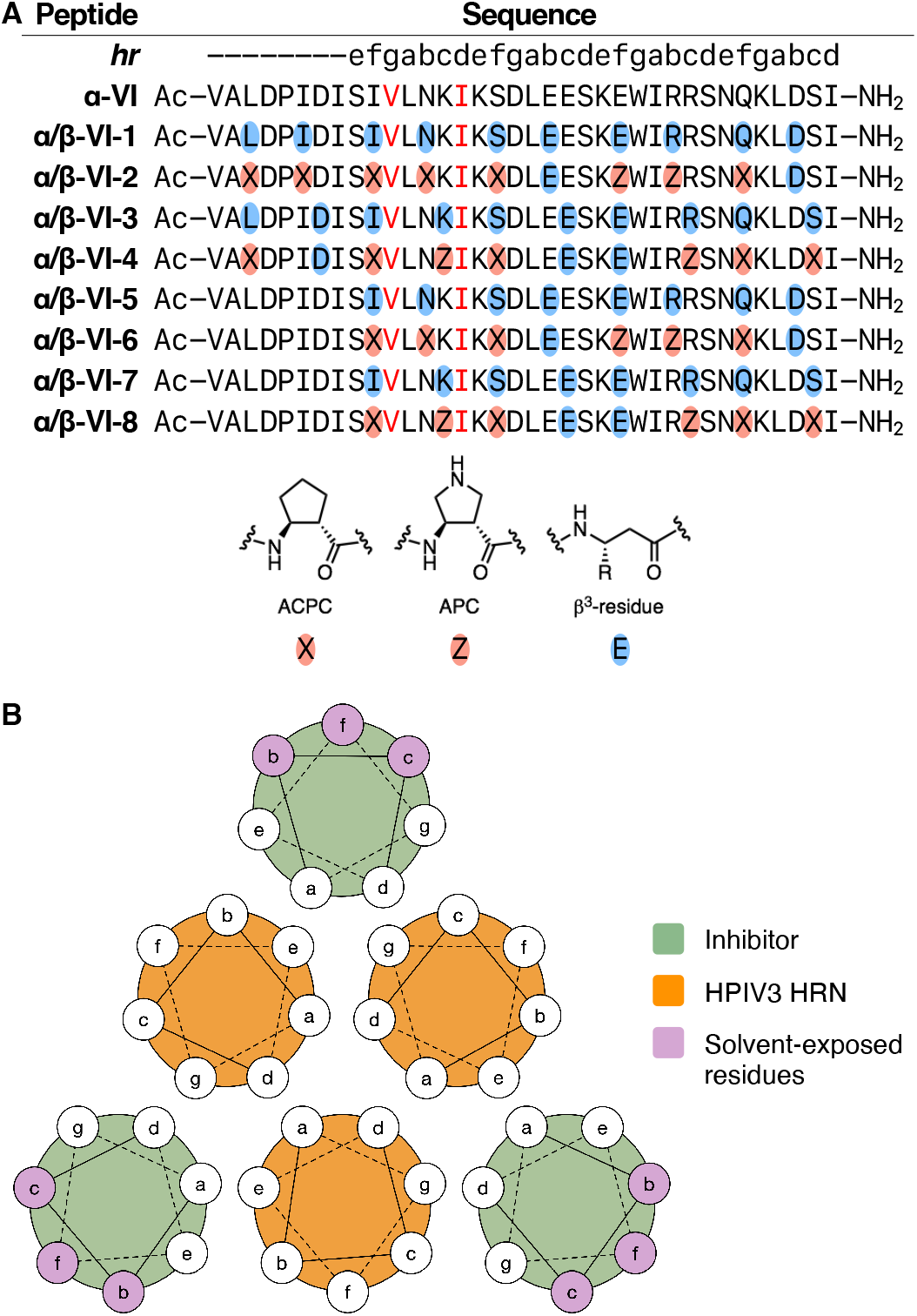
Design of α/β-peptide analogues of α-VI. (A) Sequences of α-VI and α/β-peptide variants. α-Amino acid substitutions E459V and A463I are shown in red. β3 (blue ovals) or cyclically-constrained β (red ovals) residues were incorporated either throughout the sequence (α/β-VI-1–4) or limited to the α-helical segment (α/β-VI-5–8). (B) Helical wheel diagram of an antiparallel six-helix bundle comprised of HPIV3 HRN (orange) and inhibitor (green) peptides. Inhibitor design was based on replacement of α-amino acid residues at the solvent-exposed heptad repeat positions *b*/*f* or *c*/*f* with β-residues.

Three α/β-peptide design variables were explored (Figure 1A). (1) Two different replacement registries that produce an ααβαααβ pattern were examined, one locating β residues at *b* and *f* positions (α/β-VI-1, α/β-VI-2, α/β-VI-5 and α/β-VI-6), and the other locating β residues at *c* and *f* (α/β-VI-3, α/β-VI-4, α/β-VI-7 and α/β-VI-8). Previous work has shown that α/β-peptides with a repeating ααβαααβ pattern adopt an α-helix-like conformation and display substantial resistance to proteolysis.^33-41^ (2) Two different extents of β substitution were evaluated, one with β residues throughout the entire length of α-VI (α/β-VI-1, α/β-VI-2, α/β-VI-3 and α/β-VI-4), and the other lacking β residues near the N-terminus (α/β-VI-5, α/β-VI-6, α/β-VI-7 and α/β-VI-8). The latter set was designed based on the crystal structure of the α-VI+HPIV3 HRN 6HB assembly, in which the eight N-terminal residues of α-VI adopt an extended conformation.^27^ A similar extended segment was observed in a structure of the HPIV3 F ectodomain in its post-fusion state.^42^ (3) The type of β-amino acid residues was varied. In one set of analogues (α/β-VI-1, α/β-VI-3, α/β-VI-5 and α/β-VI-7), each replacement involved the β^3^ homologue of the original α residue, which preserves the original side chain. In the second set (α/β-VI-2, α/β-VI-4, α/β-VI-6 and α/β-VI-8), many of the β residues featured a five-membered ring constraint, which stabilize the helical conformation relative to intrinsically flexible β^3^ residues.^33,34,37^

### α/β derivatives of α-VI form stable assemblies with the HRN domain of HPIV3

We investigated the interactions of α-VI and α/β-peptide variants with a peptide corresponding to the HRN domain of HPIV3 F (“HPIV3 HRN”) by circular dichroism (CD) spectroscopy. HPIV3 HRC-derived peptides, such as α-VI, co-assemble with HPIV3 HRN in solution to form six-helix bundles similar to those found in the post-fusion state of HPIV3 F.^27^ Mixing HPIV3 HRN with α/β-peptide variants of the HRC domain led to the appearance of strong CD signals that are consistent with formation of helical co-assemblies (Figure 2A).^33,36-38^

**Figure 2.**
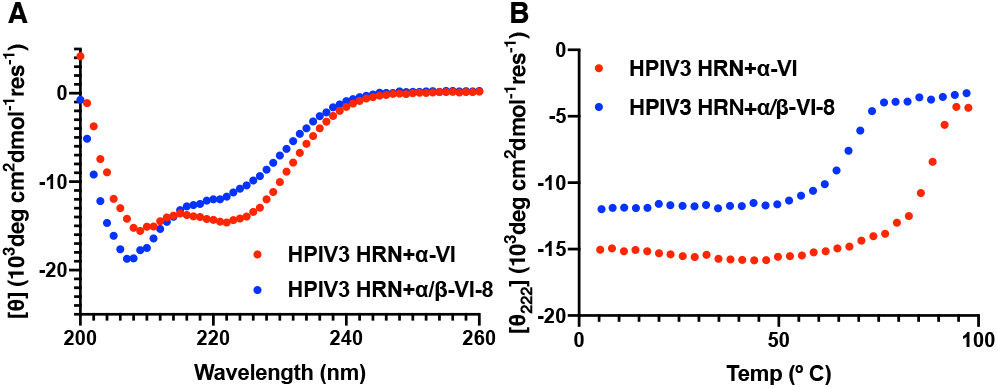
Circular dichroism studies of HPIV3 HRN + VI variant co-assemblies. (A) Circular dichroism (CD) spectra of 1:1 mixtures of α-VI (red) and α/β-VI-8 (blue) with HPIV3 HRN at 50 µM in PBS. (B) Temperature-dependent denaturation of co-assemblies formed by 1:1 mixtures of α-VI (red) and α/β-VI-8 (blue) with HPIV3 HRN at 50 µM in PBS.

Variable-temperature CD measurements can be used to assess the stability of 6HB assemblies.^27^ The co-assembly formed between HPIV3 HRN and α-VI exhibited an apparent thermal denaturation temperature (*T*_m,app_; mid-point of the denaturation process) of 88 °C (Figure 2B). All α/β-peptides formed co-assemblies with HPIV3 HRN that denatured at lower temperatures relative to the HPIV3 HRN+α-VI co-assembly (Table 1). Co-assemblies with α/β-peptides containing β residues throughout the entire sequence (α/β-VI-1, α/β-VI-2, α/β-VI-3 and α/β-VI-4) were less stable (lower *T*_m,app_) than co-assemblies of α/β-peptides lacking β residues near the N-terminus (α/β-VI-5, α/β-VI-6, α/β-VI-7 and α/β-VI-8). Among this latter group, the α/β-peptides that contained cyclic residues (α/β-VI-6 and α/β-VI-8) formed more stable assemblies than did the analogues that contained only flexible β^3^ residues (α/β-5 and α/β-7). This trend is consistent with past observations regarding the stabilizing effects of replacing β^3^ residues with cyclic β residues. Peptide α/β-VI-8, which contains six cyclically-constrained β residues and two β^3^-hGlu (β3E) residues at heptad positions *c* and *f* within the C-terminal segment, formed the most stable co-assembly with HPIV3 HRN and was selected for further characterization.

**Table 1.**
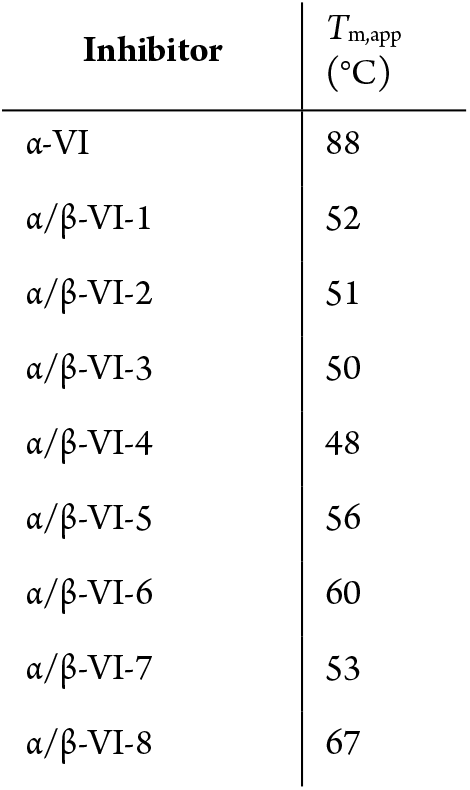
**Apparent melting temperatures (*T***_***m***,***app***_**) of co-assemblies formed between VI variants and HPIV3 HRN**

### Crystal structure of α/β-VI-8 co-assembled with the HRN domain of HPIV3 F

The crystal structure of α/β-VI-8 + HPIV3 HRN was determined (Figure 3B) and shows a 6HB assembly similar to that observed within the HPIV3 F protein in the post-fusion conformation.^42^ Three copies of α/β-VI-8 are arrayed around the core formed by three HRN peptides. Each α/β-VI-8 molecule displays an α-helix-like conformation spanning residues 457–484 (numbering based on the HPIV3 F sequence); each turn of this helix, however, contains one more carbon atom relative to an authentic α-helical turn. The extended conformation of the N-terminal eight residues of α/β-VI-8 matches the corresponding segment of full-length F in the post-fusion state^42^ and of α-VI + HPIV3 HRN.^27^ All eight β residues of α/β-VI-8 align along one side of the helix, and none of the β residue side chains contacts the HRN core (Figure 3C-F). Thus, the α/β-VI-8 + HPIV3 HRN co-crystal structure shows that our design goal of sequestering β residues away from the HRC-HRN interface was achieved.

**Figure 3.**
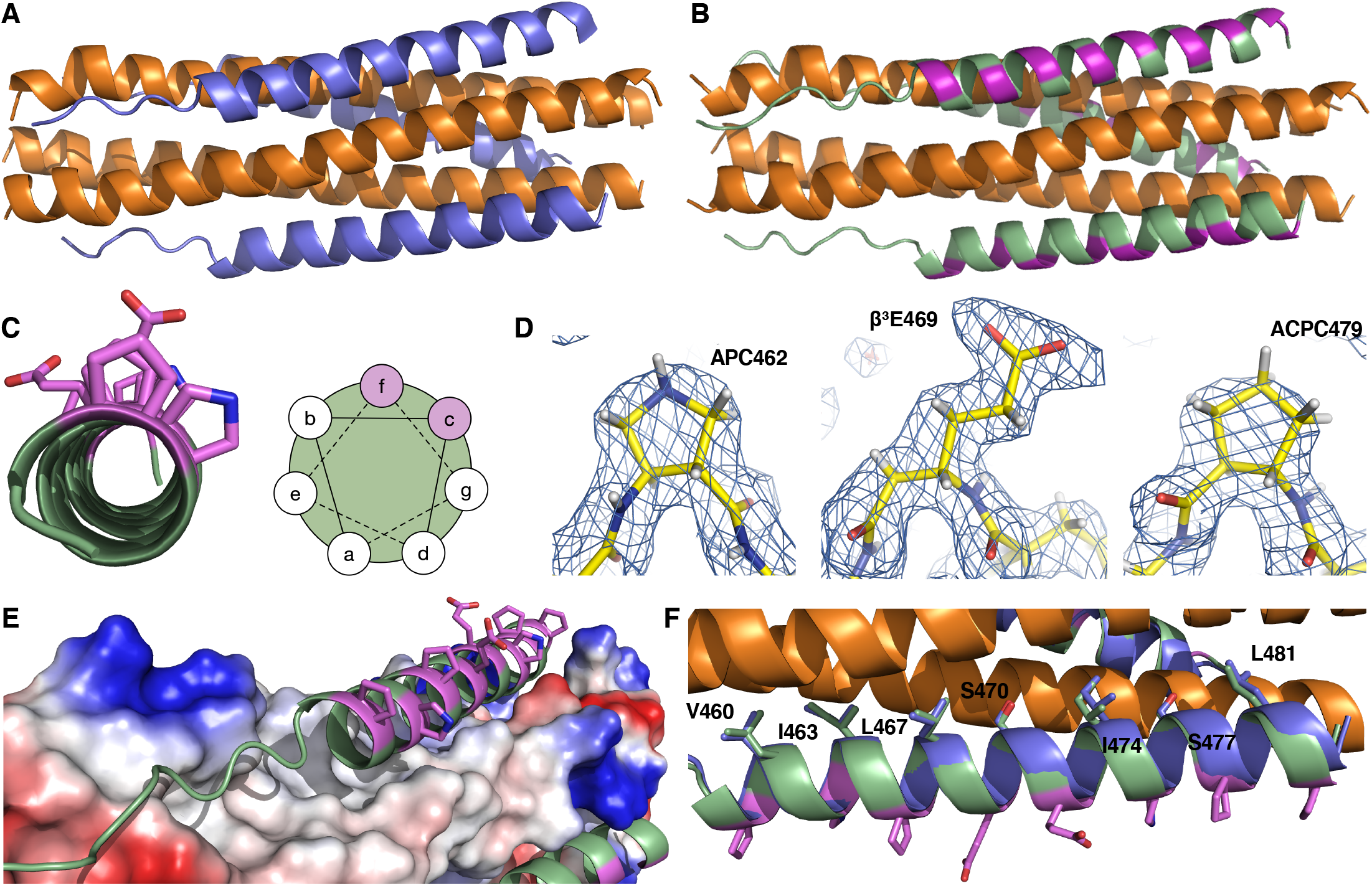
Crystal structure of α/β-VI-8 bound to the HRN domain of HPIV3. (A) The 6HB co-assembly formed by HPIV3 HRN (orange) and α-VI (blue) (PDB 6NYX). (B) The 6HB co-assembly formed by HPIV3 (orange) and α/β-VI-8 (green, β residues depicted as violet) (PDB 6VJO). (C) Helix-wheel diagram showing orientation of β residues (violet) at *c* and *f* positions of the heptad repeat as designed. (D) Electron-density maps of representative β residues APC462 (left), β_3_E469 (middle), and ACPC479 (right). (E) α/β-VI-8 binds within a hydrophobic groove on the HPIV3 HRN trimer (shown as map of negative (red), neutral (gray), or positive (blue) electrostatic potential. (F) Overlay of α-VI (blue) and α/β-VI-8 (green, β residues depicted as violet). Conformations of interfacial amino acid residues were conserved.

### Inhibition of HPIV3 F-mediated cell-cell fusion by α/β analogues of α-VI

We evaluated inhibition of fusion mediated by HPIV3 F co-expressed with HPIV3 HN using a cell–cell fusion assay based on β-galactosidase complementation (Supplemental Figure 1).^43,44^ While we anticipated that maximum inhibitory potency in this assay would require lipid conjugation of HRC-based peptides, in preliminary studies we compared the eight α/β-peptide analogues of α-VI (Figure 1A) without lipid modification. The trends for cell-cell fusion inhibitory potency were similar to those observed in the variable-temperature CD studies. The α/β-peptides containing β residues throughout the entire sequence failed to block cell-cell fusion (not shown). Among the four α/β-peptides lacking β residues in the N-terminal segment, the two containing cyclic residues (α/β-VI-6 and α/β-VI-8) were the most potent inhibitors. Neither, however, matched α-VI as an inhibitor of F-mediated fusion. The α/β-peptides containing only β^3^ residues (α/β-VI-5 and α/β-VI-7) were less effective than the analogues containing cyclic β residues. We focused on α/β-VI-8 for subsequent studies involving lipid-conjugated peptides.

We evaluated derivatives of α-VI and α/β-VI-8 bearing a C-terminal cholesterol appendage (Figure 4A) as inhibitors of F-mediated cell-cell fusion.^45^ Each 36-residue sequence was extended at the C-terminus by the segment Gly-Ser-Gly-Ser-Gly-Cys. Reaction of the Cys thiol with a bromoacetyl derivative of cholesterol containing a 4-unit PEG spacer (PEG_4_) generated the lipopeptides. The PEG_4_ segment provides a flexible linker between the peptide inhibitor and the membrane-embedded cholesterol moiety and enhances inhibitory activity in HPIV3 HRC-derived peptides.^23,25^ The resulting molecules were designated “α-VI–PEG_4_–Chol” and “α/β-VI-8–PEG_4_– Chol”.

**Figure 4.**
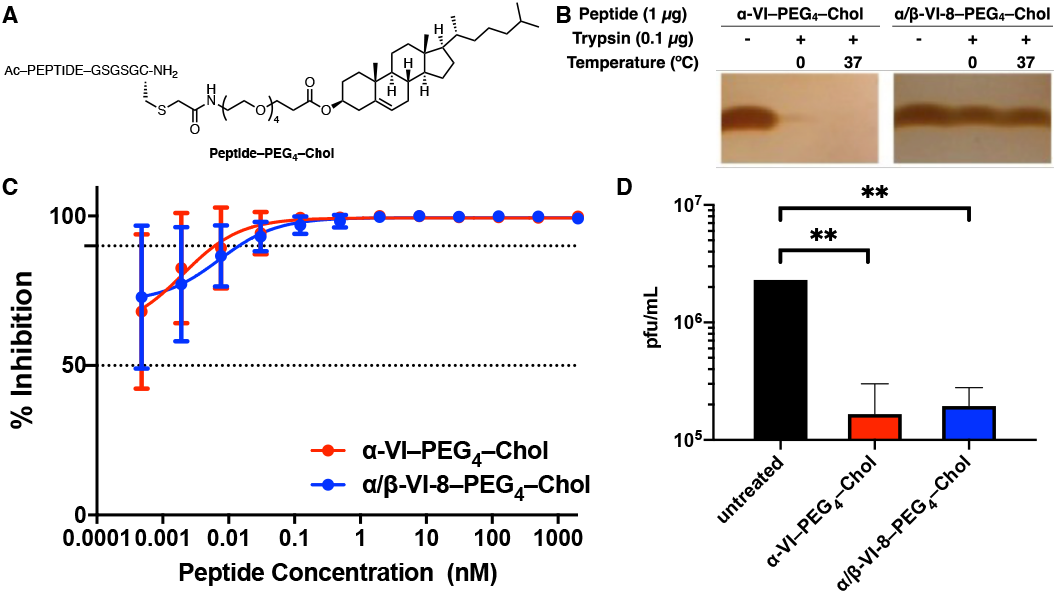
Proteolytic resistance and antiviral efficacy of α/β-VI-8– PEG_4_–Chol. (A) Structure of cholesterol-conjugated peptides. (B) Degradation of α-VI–PEG_4_–Chol (left) and α/β-VI-8–PEG_4_–Chol (right) by incubation with trypsin at 0 and 37 °C. (C) Inhibition of HPIV3 infection in cell monolayers by α-VI–PEG_4_–Chol (red) or α/β-VI-8– PEG_4_–Chol (blue). Percent inhibition was calculated as the ratio of plaque forming units in the presence of a specific concentration of inhibitor and the plaque forming units in the absence of inhibitor. Each point represents the mean from three separate experiments ± SEM with the curve representing a three-parameter dose-response model. (D) Inhibition of HPIV3 infection in human airway epithelial (HAE) cultures by α-VI–PEG_4_–Chol (red) or α/β-VI-8–PEG_4_–Chol (blue). HAE cultures were treated with α-VI–PEG_4_–Chol or α/β-VI-8–PEG_4_–Chol at 1 μM and infected with 5000 pfu/well of hPIV3 CI-eGFP. After 2 h, the peptides were removed, and supernatants were collected 1 day post infection. Infectious viruses released were quantified by titration. Data are depicted as the mean from three separate experiments ± SD. Data were analyzed by Student’s two-tailed unpaired *t* test (\*\**P* < 0.01).

### α/β-VI-8-PEG_4_-Chol exhibits enhanced resistance to protease-mediated degradation

The specific sites at which α-VI and α/β-VI-8 may be proteolytically cleaved in vivo are unknown; we evaluated relative degradation in the presence of the aggressive pro-tease trypsin to compare α-VI–PEG_4_–Chol and α/β-VI-8–PEG_4_– Chol (Figure 4B). The cholesterol-conjugated derivative of α/β-VI-8 was stable under conditions that rapidly caused degradation of the analogous α-lipopeptide. After one-hour incubation with trypsin at 0 °C, nearly all α-VI–PEG_4_–Chol was degraded. After incubation with trypsin at 37 °C, α-VI–PEG_4_–Chol could not be detected. However, α/β-VI-8–PEG_4_–Chol remained intact under these conditions, which demonstrates the substantial inhibition of proteolysis provided by periodic α-to-β replacement.

### α/β-VI-8-PEG_4_-Chol potently inhibits HPIV3 infection in monolayer cultured cells

We assessed the efficacy of α/β-VI-8– PEG_4_–Chol at inhibiting infection by HPIV3 in cell culture (Figure 4C). We used a well-characterized HPIV3 clinical isolate virus (“CI”) because we have shown that the HN-F glycoprotein fusion complexes of circulating viruses differ significantly from the HN and F of standard laboratory-grown viruses.^46-51^ The fusion complex in laboratory culture-adapted viruses is more readily activated to promote fusion and less sensitive to inhibition by HRC peptides compared to the fusion complex in authentic circulating HPIV3, making it essential to assess inhibitors using authentic circulating HPIV3.^46-51^ α/β-VI-8–PEG_4_–Chol potently inhibited infection by HPIV3 CI, with potency similar to that of α-VI–PEG_4_–Chol. Both cholesterol-conjugated peptides achieved 100% inhibition at 0.1 nM (Fig. 4C), while the corresponding unconjugated peptides displayed IC_50_ for fusion inhibition of ∼10 nM. This dramatic difference in potency between unconjugated and lipid-conjugated peptides is consistent with previous observations.^23-26^

### α/β-VI-8-PEG_4_-Chol potently inhibits HPIV3 infection in human airway epithelium

Human airway epithelium (HAE) cultures have proven to be an authentic model of human lung, reflecting the cell environment and selective pressure of the natural tissue^46,48^, and the efficacy with which fusion inhibitors block viral infection in HAE seems to correlate with in vivo efficacy.^23,46^ We compared the antiviral effects of α-VI–PEG_4_–Chol and α/β-VI-8–PEG_4_–Chol in HAE. A single dose of 1 µM (final concentration) of either α-VI– PEG_4_–Chol or α/β-VI-8–PEG_4_–Chol decreased the viral titer of HPIV3 CI-eGFP by 98% at day 1 (Figure 4D). At day 2, α-VI– PEG_4_–Chol treatment reduced titer by 80% while α/β-VI-8–PEG_4_– Chol reduced titer by close to 98% (data not shown).

### Biodistribution: α/β-VI-8-PEG_4_-Chol reaches higher concentrations than does α-VI-PEG_4_-Chol in serum and lung tissue

The utility of a fusion-inhibitory peptide as an antiviral agent depends on availability at the site of infection. We assessed biodistribution of α-VI–PEG_4_–Chol and α/β-VI-8-PEG_4_-Chol in cotton rats after either intranasal or subcutaneous administration. α/β-VI-8– PEG_4_–Chol was present at higher concentration than was α-VI– PEG_4_–Chol in the lung 24 hours after either intranasal or subcutaneous administration (Figure 5A). A substantially higher concentration of α/β-VI-8–PEG_4_–Chol was achieved in the lung upon intranasal administration relative to subcutaneous administration. The α/β-lipopeptide also demonstrated a longer serum half-life than the α-lipopeptide after either administration route (Figure 5B). The concentration difference in the lung, particularly after intranasal administration, indicates a major advantage for the backbone-modified peptide, since persistence at the relevant tissue site should improve efficacy *in vivo*.

**Figure 5.**
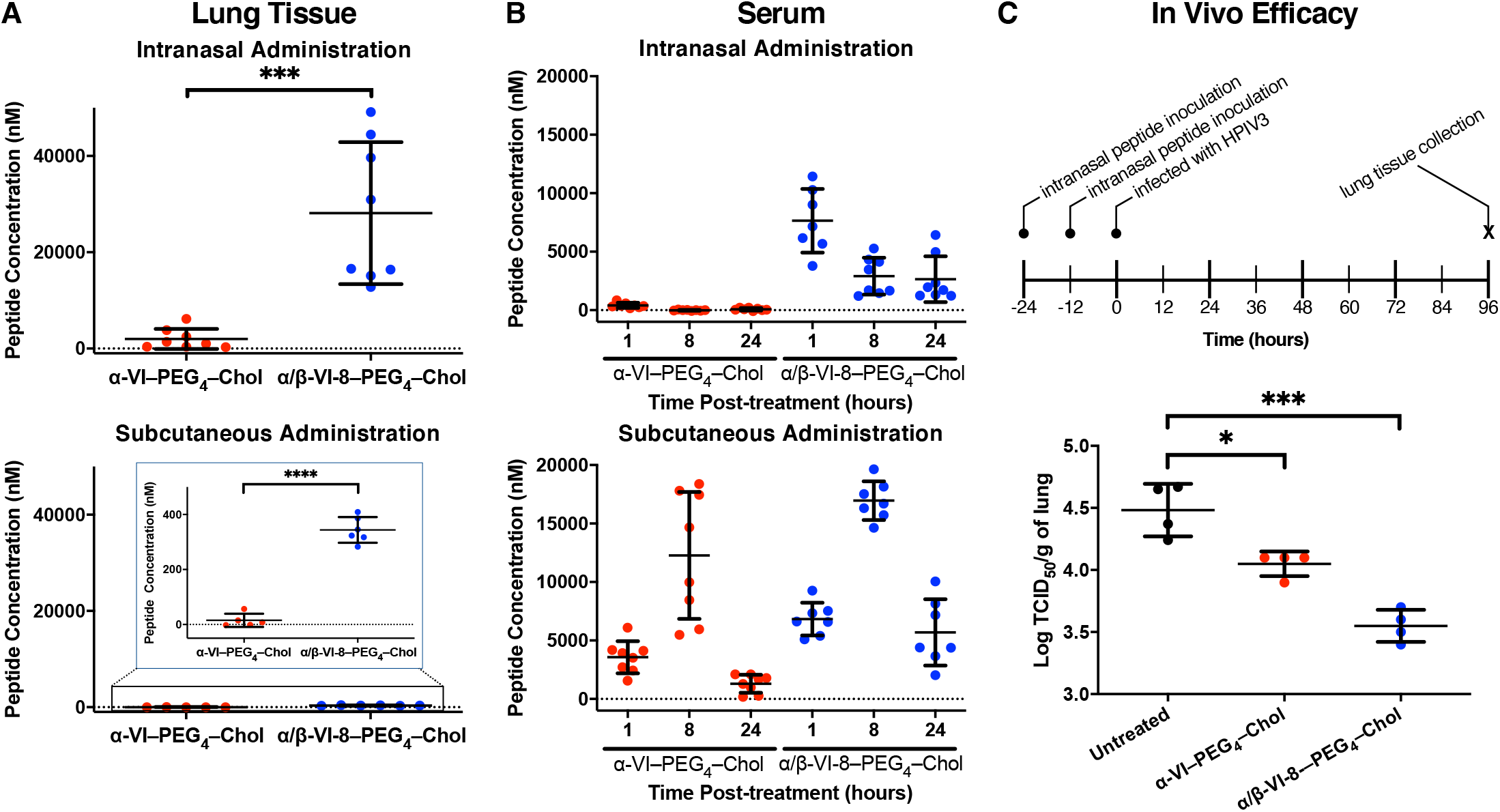
Biodistribution and efficacy of α/β-VI-8–PEG_4_–Chol *in vivo*. (A) Biodistribution of α/β-VI-8–PEG_4_–Chol in cotton rat lung tissue. Peptide concentration in the lung was determined by sandwich ELISA 24 h after intranasal (top) or subcutaneous (bottom) administration of lipopeptide. Data are depicted as the mean from eight (intranasal) or six (subcutaneous) separate experiments ± SD. Data were analyzed by Student’s two-tailed unpaired *t* test (\*\*\**P* < 0.001 and \*\*\*\**P* < 0.0001). (B) Biodistribution of α/β-VI-8–PEG_4_–Chol in serum. Peptide concentration in serum was determined by sandwich ELISA 24 h after intranasal (top) or subcutaneous (bottom) administration of lipopeptide. Data are depicted as the mean from eight separate experiments ± SD. (C) *In vivo* efficacy of α/β-VI-8–PEG_4_–Chol against HPIV3 infection in cotton rats with (top) schematic of experimental design and (bottom) data depicted as the mean from four separate animals ± SD. Data were analyzed by Tukey’s multiple comparison test (\*\**P* < 0.01, \*\*\*\**P* < 0.0001).

### Inhibition of HPIV3 infection *in vivo* by α/β-VI-8-PEG_4_-Chol

To assess *in vivo* efficacy of the α/β-lipopeptide, cotton rats were administered 2 mg/kg of either α-VI–PEG_4_–Chol or α/β-VI-8-PEG_4_-Chol peptides intranasally, at 24 and 12 hours before infection and then infected with 10^6^ pfu of HPIV3 CI; the experimental scheme is shown in Figure 5C (top). Three days after infection, lung viral titers were measured. Figure 5C (bottom) shows viral titer in lipopeptide-treated animals compared to untreated animals. α/β-VI-8-PEG_4_-Chol was superior to α-VI-PEG_4_-Chol, with the α/β-lipopeptide reducing viral titer by over 10-fold relative to no treatment (*P* < 0.0001using Tukey’s multiple comparison test. α/β-VI-8-PEG_4_-Chol reduced viral titer by 0.5 log more than the α-VI-PEG_4_-Chol (*P* < 0.01). When the lower concentration of 0.4 mg/kg of peptide was administered 24 and 12 hours before infection, a less than 10-fold but nonetheless significant reduction in viral titer was achieved (Supplementary Figure 2). A comparable reduction in humans would be clinically significant.^52^

## Discussion

The results presented above suggest that modification of an HRC-derived peptide via periodic α-to-β replacement at positions selected to avoid the HRC-HRN interface in the 6HB assembly results in maintenance of fusion-inhibitory activity and inhibition of viral entry, while hindering proteolytic degradation. The resistance to proteolysis leads to enhanced persistence and antiviral performance *in vivo* for an α/β-lipopeptide relative to a comparable α-lipopeptide. This study represents, to our knowledge, the first *in* vivo evaluation of an α/β-peptide inhibitor of viral infection and lays the foundation for a new type of agent to address HPIV3 infections in a prophylactic and/or therapeutic manner.

Because entry is the essential first step of infection by enveloped viruses, the inhibition of fusion protein structural transitions is an attractive basis for antiviral therapy.^8^ The advance we have achieved in developing α/β-VI-8–PEG_4_–Chol builds upon earlier work with α-lipopeptide entry inhibitors in which we showed that specific features of amino acid sequence, identity of the lipid moiety, linker length, and solubility are all important for anti-viral efficacy.^20-27^ Our observations suggest that further improvements in α/β-peptide activity may be possible by pursuing two strategies. (1) Variable-temperature CD data suggest that the 6HB assembly formed by α/β-VI-8 with the HPIV3 HRN is less stable than the 6HB assembly formed by α-VI and the HPIV3 HRN. We predict that modifications to α/β-VI-8 that lead to enhanced 6HB stability will result in improved antiviral potency. (2) Although α/β-VI-8–PEG_4_–Chol is highly resistant to degradation by trypsin, it is possible that the N-terminal segment of α residues in this α/β-lipopeptide or the C-terminal α residue segment used to connect the cholesterol moiety is susceptible to cleavage by other proteases in the lung. Incorporation of one or two β residues in these segments could therefore improve lipopeptide persistence *in vivo*. In this regard, we note recent results indicating that several positions in the N-terminal segment of an HRC peptide related to α-VI tolerate α-to-β substitution without loss of potency for inhibiting HPIV3 infection.^53^

The favorable performance of our lead α/β-lipopeptide in the HAE system (Figure 4D) is promising in terms of our long-term aim to develop backbone-modified peptides for clinical use. We previously showed that the fusion complex formed by HN and F for HPIV3 differs in important ways between clinical isolate (CI) viruses (*i*.*e*., those that can grow in humans) and laboratory-adapted strains.^46-51^ Viruses bearing the fusion complex of clinical strains grow efficiently in HAE and *in vivo*, but fail to grow on immortalized cells because of the profound specificity of the CI viral fusion machinery for the authentic host.^46-48,54^ We posit that it is critical to assess antiviral agents, such as those introduced here, in the context of the viral fusion complexes that exist in human infections (*i*.*e*., to assess efficacy of inhibiting viral infection in authentic lung tissues).

In the cotton rat experiment (Figure 5C), the superiority of α/β-VI-8-PEG_4_-Chol over the α-lipopeptide, α-VI-PEG_4_-Chol, must be attributed to the resistance of the α/β-lipopeptide to proteolysis (Figure 4B), which we propose to be the source of the significantly enhanced persistence of the α/β-lipopeptide relative to the α-lipopeptide *in vivo* (Figure 5A-B). The efficacy of the α/β-lipopeptide *in vivo* is especially striking given that the six-helix bundle assembly formed by α/β-VI-8 with HPIV3 HRN is significantly less stable than the assembly formed by α-VI, and α/β-VI-8 is less effective in inhibiting F-mediated cell-cell fusion than is α-VI (Supplementary Figure 1).

Two major factors augment the potential for clinical application of our α/β-lipopeptide inhibitors. First, the ability to administer these inhibitors intranasally offers a tractable and practical route of delivery. In fact we recently showed for another respiratory virus (SARS-CoV-2) that daily intranasal peptide administration completely blocked infection of naïve animals by infected animals.^55^ These results lend support to the feasibility of using intranasal anti-HPIV3 peptides to prevent infection of vulnerable individuals (*e*.*g*., during immune compromise, stem cell or organ transplant, etc.). Second, α/β-lipopeptides are chemically stable and should tolerate storage at room temperature for long periods, which means that no cold chain would be required. These factors suggest α/β-lipopeptides could be useful even outside of advanced health care settings,

## Materials and Methods

### Cells

293T (human kidney epithelial), CV-1 and Vero E6 cells were obtained from ATCC and were grown in Dulbecco’s modified Eagle’s medium (DMEM) (Gibco) supplemented with 10% fetal bovine serum and antibiotics at 37 °C and 5% CO_2_. All cells tested negative for mycoplasma (MycoAlert™ Mycoplasma Detection Kit (Lonza)).

### Peptide synthesis

All peptides were produced by standard Fmoc-based solid-phase methods. The cholesterol moiety was attached to the peptide via chemoselective reaction between the thiol group of an extra cysteine residue, added C-terminally to the sequence, and a bromoacetyl derivative of cholesterol, as previously described.^23,24,56^

### Structure Determination

Co-crystallization screening was performed using a 1:1 mixture of α/β-VI-8 and HPIV3 HRN. Crystal growth was promoted by vapor diffusion using hanging drop methods. Conditions that generated crystal hits were optimized for concentrations of peptide, salt, and polyethylene glycol (PEG), as well as pH, until crystals of sufficient quality for x-ray analysis were obtained. The optimized crystals diffracted to 2.0 Å. The structure was solved by molecular replacement using a truncated dimer of HPIV3 HRN+VIQKI (PDB: 6NRO) as a search model.^27^ Specifically, chains A and B from 6NRO were truncated to include residues 153– 173 (chain A; numbers correspond to full-length F) and 460–480 which is important given the global distribution of HPIV3 and the impact of this pathogen in developing societies. (chain B), and all hydrogen atoms and side chains were removed prior to molecular replacement.

### Viruses

HPIV3 clinical isolate virus (CI) was obtained from the Clinical Microbiology Laboratories at New York Presbyterian Hospital and grown in human airway epithelium (HAE) at an air-liquid interface for only one passage prior to use in these experiments. Recombinant CI virus used in the cotton rat experiments was generated as described.^48^

### Antiviral activity against live HPIV3

Peptide activity against HPIV3 was determined by plaque reduction assays in infected cell monolayers.^46^ Data were expressed as mean ± standard deviation (s.d.) (*n*=3 separate experiments).

### HAE cultures

The EpiAirway AIR-100 system (MatTek Corporation) consists of normal human-derived tracheo/bronchial epithelial cells that have been cultured to form a pseudostratified, highly differentiated mucociliary epithelium closely resembling that of epithelial tissue *in vivo*. Upon receipt from the manufacturer, HAE cultures were transferred to 6-well plates (containing 0.9 mL medium per well) with the apical surface remaining exposed to air and incubated at 37 °C in 5% CO_2_.

### Peptide efficacy assessment in HAE cultures

HAE cultures were infected by applying 100 µL of EpiAirway medium containing 5,000 plaque forming units (PFU) of HPIV3 with or without peptide (1 μM final concentration) to the apical surface for 90 min at 37 °C. The medium containing the inoculum was removed, and cultures were placed at 37 °C and fed each day with 0.9 mL medium via the basolateral surface. Viruses were harvested by adding 200 µL medium per well to the HAE cultures’ apical surface and allowing the cultures to equilibrate for 30 min at 37 °C. The suspension was then collected, and viral titers were determined as previously described.^46^ Viral collection was performed sequentially with the same wells of cells on each day post infection.

### Protease sensitivity of HPIV3 derived peptides

For the trypsin digestion, 1 μg of each peptide was treated with the indicated amount of trypsin in 10 µL of PBS. Peptide solutions were then incubated at 0 °C or 37 °C for 1 hour. Following incubation, 10 µL of Laemmli’s SDS reducing buffer was added to each solution. Samples were boiled for 10 min at 99 °C then run on a 4-20% Tris Glycine gel at 120 V. The gel was allowed to fix overnight in 0.0125% glutaraldehyde in PBS. Gels were stained using a Pierce® Silver Stain kit (Cat# 24600).

### Infection of cotton rats, peptide treatment, and virus titer determination

Inbred cotton rats (*Sigmodon hispidus*) were purchased from Envigo, Inc. (Indianapolis). Both female and male cotton rats at the age of 5 to 7 weeks were used for this study. Group size was based on power analysis to determine a difference in treatment of at least one log10 in viral titer after treatment with the α-VI peptide.

For intranasal (i.n.) infection, 10^6^ TCID_50_ of recombinant CI HPIV3 ^48^ was inoculated in phosphate-buffered saline to isofluorane anesthetised cotton rats in a volume of 100 µL i.n. as previously described ^46,57^. To evaluate the effect of HRC peptides, animals were inoculated i.n. with peptide (2 mg/kg or 0.4 mg/kg in 100 µL of water) 24 h and 12 h before infection. Four days after infection, the animals were asphyxiated using CO_2_, and their lungs were collected and weighed. Lung tissue was minced with scissors and dounced with a glass homogenizer. Serial 10-fold dilutions of supernatant fluids were assessed for the presence of infectious virus in 48-well plates as described.^48^ The TCID_50_was calculated as described previously. The animal experiments were approved by the Institutional Animal Care and Use Committee of The Ohio State University.

### Biodistribution study methods

#### Animal experiment

For biodistribution experiments in cotton rats, the animals received the indicated peptides (2 mg/kg) either i.n. of s.q. in 100 µL of diluent. At 1 h and 8 h post administration, blood was collected retroorbitally in EDTA tubes from all animals, centrifuged for 10 min at 2000 RPM, and plasma was transferred into new tubes and stored at −20°C for use in ELISA. At 24 h after administration, the animals were euthanized, blood was collected by intracardiac puncture in EDTA vacutainer tubes, and sera were conserved at -20 °C until their use in ELISA. Organs from each animal were collected and conserved at - 80 °C. The tissues were separated in two: one half frozen on dry ice and conserved at −80°C for ELISA.

#### Enzyme-linked immunosorbent assay (ELISA) for determination of peptide concentration

ELISA was performed from frozen samples as previously described.^28^ Each organ was weighed and mixed in PBS (1:1, w/vol) using an ultra turrax homogenizer. Samples were then treated with acetonitrile/1% TFA (1:4, vol/vol) for 1 h on a rotor at 4 °C and then centrifuged for 10min at 8000 rpm. Maxisorp 96 well plates (Nunc) were coated overnight with 20 µg/mL affinity-purified Ab (anti α-VI and anti α/β-VI-8 (custom made by Gene-Script)) in carbonate/bicarbonate buffer (pH 7.4) at 4 °C. Plates were washed 3x with PBS, then blocked with PBS / 3% BSA for 2 hours and placed on ice. Serial dilutions of organ and serum samples were prepared, and the blocking buffer was replaced by the serial dilutions of each sample added in 3% PBS-BSA (pH 7.4). The samples were incubated at RT for 2.5 hours. The plates were washed 4x with 150 µL of PBST (0.05%), and 100 µL / well of purified rabbit biotin-conjugated and biotin-conjugated anti-α/β-VI-8 (horse radish peroxidase [HRP]-conjugated) at 1:500 in PBS BSA (3%) were added to the plates and incubated at RT for 2 h. After washing the plates 4x with PBST and 2x with PBS, streptavidin HRP (1:1000 in PBS) was added, and the plates were incubated for 30 min at RT, washed 4x with PBST, and 3x with PBS before addition of 1-Step^TM^ Ultra TMB-ELISA (ThermoFisher #34028). The plates were covered and monitored for development, 50 µL of 2 M sulfuric acid was added to each well to stop the reaction, and results were recorded as absorbance at 450 nm on the Infinity PRO1000 (Tecan). Standard curves were established for each peptide using 1:2 serial dilutions from 40 nM to 0 nM (using the same ELISA conditions as for the test samples), and the detection limit was determined to be 0.15 nM.

### Statistical Analysis

Data are described as mean±s.d. unless otherwise stated. The statistical analysis performed on *in vivo* experiments for comparison of HPIV3 viral titers in cotton rat lungs were based on Tukey’s multiple comparison test. All the data and statistical tests were performed in Prism 5 (GraphPad Software, La Jolla, CA).

## ASSOCIATED CONTENT

### Supporting Information

Supporting Information is available free of charge on the ACS Publications website at DOI:

Additional figures, method descriptions, and characterization data for all synthesized peptides (PDF)

## AUTHOR INFORMATION

## Author Contributions

## ACKNOWLEDGMENT

This work was supported by NIH grants R01AI114736 to A.M and R01 GM056414 to S.H.G., and by the Sharon Golub fund. V.K.O. was supported in part by NIH F32 GM122263. R.W.C. was supported in part by NIH T32 GM008349.

---

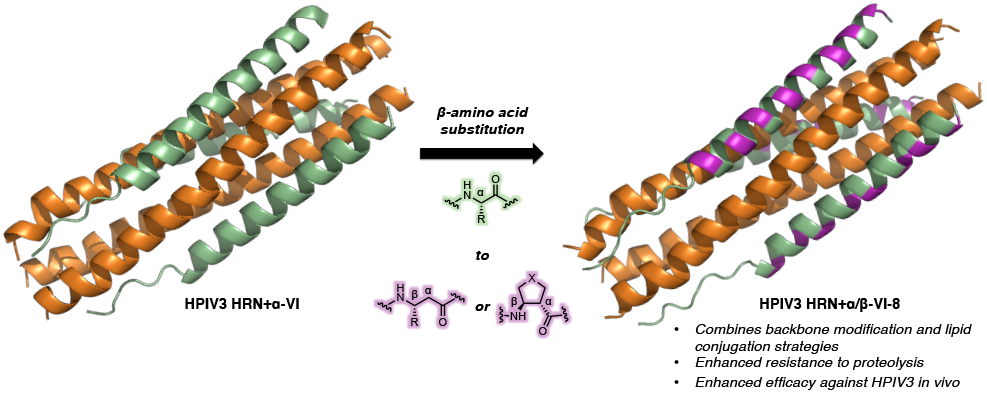

Insert Table of Contents artwork here

