## Supporting Information for "Engineering protease-resistant peptides to inhibit human parainfluenza viral respiratory infection"

#### Supplemental Figures

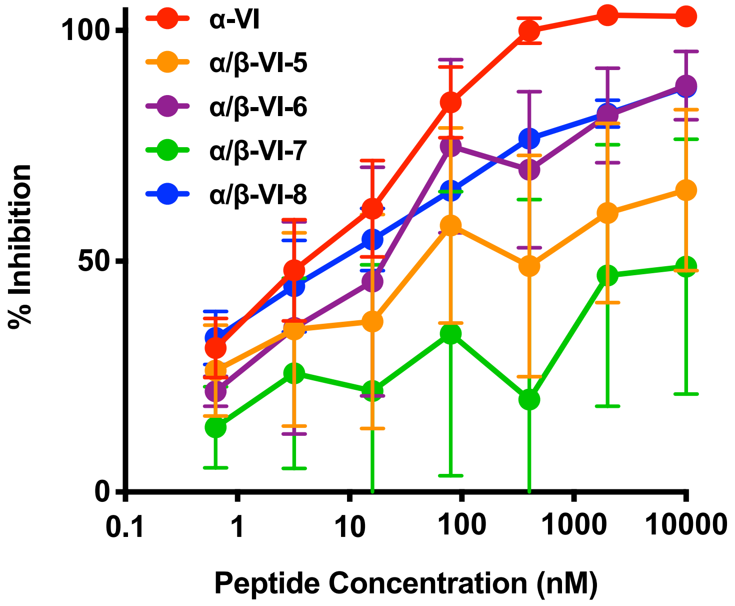

**Supplementary Figure 1. Inhibition of cell-cell fusion mediated by HPIV3 F in 293T cells by α-VI and α/β-VI variants.** Percent inhibition was calculated as the ratio of relative luminescence units in the presence of a specific concentration of inhibitor and the relative luminescence units in the absence of inhibitor and corrected for background luminescence. Each point represents the mean from three separate experiments ± SEM with lines connecting adjacent points.

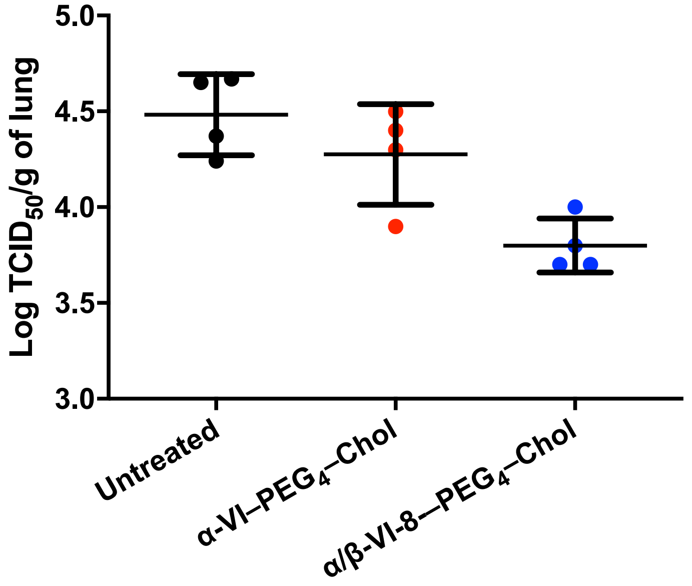

**Supplementary Figure 2.** *In vivo* efficacy of α/β-VI-8–PEG$_4$–Chol against HPIV3 infection in cotton rats. Data are depicted as the mean from four separate animals ± SD.

### Peptide Synthesis and Purification

#### Instrumentation

Solid-phase peptide synthesis was performed on a CEM MARS microwave reactor using polypropylene syringes fitted with a porous disc (Torviq). Preparative HPLC was performed using a Shimadzu HPLC system (SCL-10VP system controller, LC-6AD pumps, SIL-10ADVP autosampler, SPD-10VP UV-vis detector, FRC-10A fraction collector) equipped with a Waters XSelect CSH Prep C18 column (5 µm particle size, 19 mm × 250 mm). Peptide purity measurements were performed on a Waters Acquity H-Class UPLC equipped with an Acquity UPLC BEH C18 column (130 Å pore size, 1.7 µm particle size, 2.1 mm × 50 mm). Mass spectra were obtained on a Bruker microflex LRF MALDI-TOF-MS. Circular dichroism experiments were performed on an Aviv Biomedical model 420 CD spectrometer.

| Instrument Name | Instrument Type | Grant |
|---|---|---|
| Waters Acquity H-Class | UPLC | DARPA N66001-15-2-4023 |
| Bruker microflex LRF | MALDI-TOF-MS | Generous gift from the Bender Fund |
| Aviv Biomedical model 420 | CD spectrometer | NIH R01GM056414 |

#### General Procedure

Peptides were prepared on Rink amide resin using microwave-assisted solid-phase peptide synthesis (MA-SPPS). Resin was purchased from Millipore–Sigma. Fmoc-amino acids and coupling reagents were purchased from Chem-Impex International. Protected Fmoc-α-amino acids included: Asp(*t*-Bu ester), Glu(*t*-Bu ester), His(trityl), Lys(Boc), Asn(trityl), Gln(trityl), Arg(Pbf), Ser(*t*-Bu ether), Thr(*t*-Bu ether), Trp(Boc), and Tyr(*t*-Bu ether).

Rink amide resin was pre-swelled with DMF in a polypropylene fritted syringe, then drained and washed with DMF. Coupling reactions were performed using solutions comprised of 4 equivalents Fmoc-amino acid, 4 equivalents of 1-[bis(dimethylamino)methylene]-1H-1,2,3-triazolo[4,5-b]pyridinium 3-oxide hexafluorophosphate (HATU), and 8 equivalents of diisopropylethylamine (DIEA) in biotechnology-grade dimethylformamide (DMF) at a final concentration of 100 mM Fmoc-amino acid. Coupling reactions were carried out by microwave-assisted synthesis using a 2 min ramp to 70 °C followed by a 4 min hold at 70 °C. Deprotection was effected by addition of 20% (v/v) piperidine in biotechnology-grade DMF. The deprotection reactions were carried out by microwave-assisted synthesis using a 2 min ramp to 80 °C followed by a 2 min hold at 80 °C. The resin was washed with 3–5 resin volumes of biotechnology-grade DMF after each coupling and deprotection reaction.

**α-VI**

Sequence: Ac-VALDPIDISIVLNKIKSDLEESKEWIRRSNQKLDSI-$NH_2$

Calculated monoisotopic [M+H]: 4206.3

Calculated monoisotopic [M+2H]: 2103.7

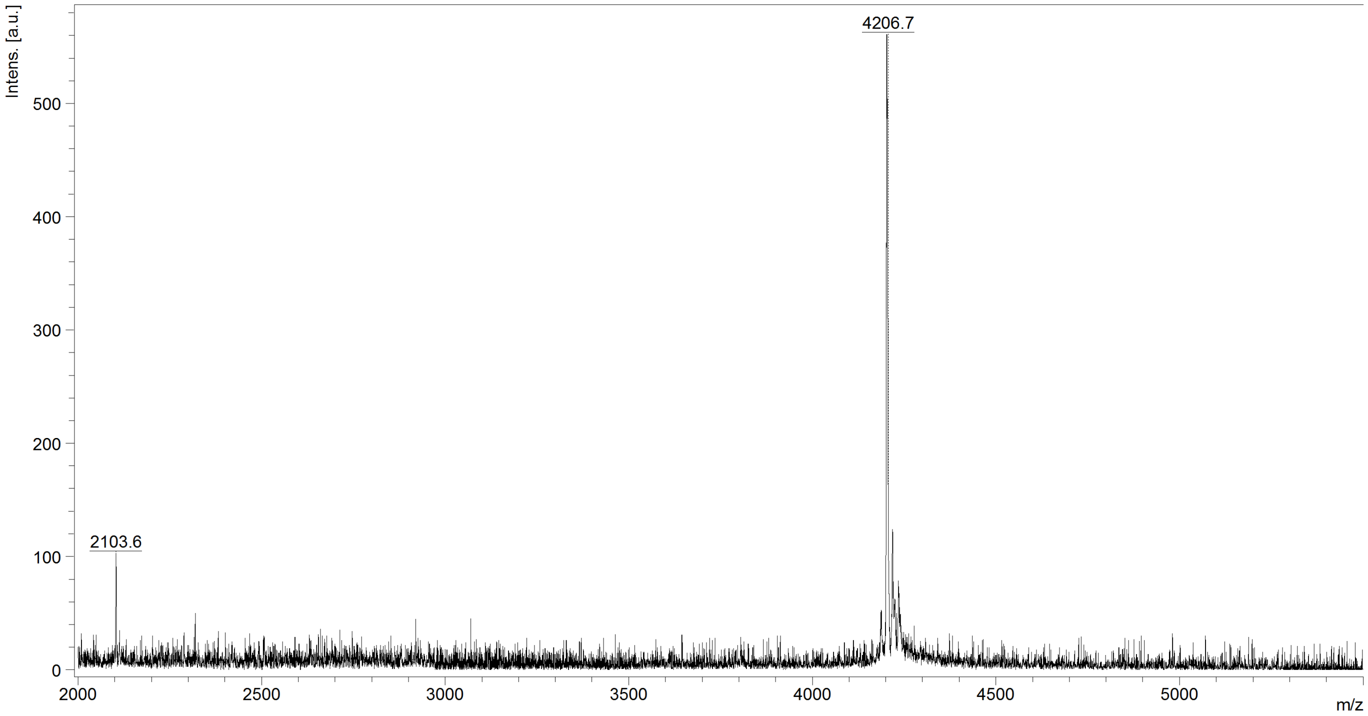

Supplementary Figure 3. α-VI MALDI-TOF analysis

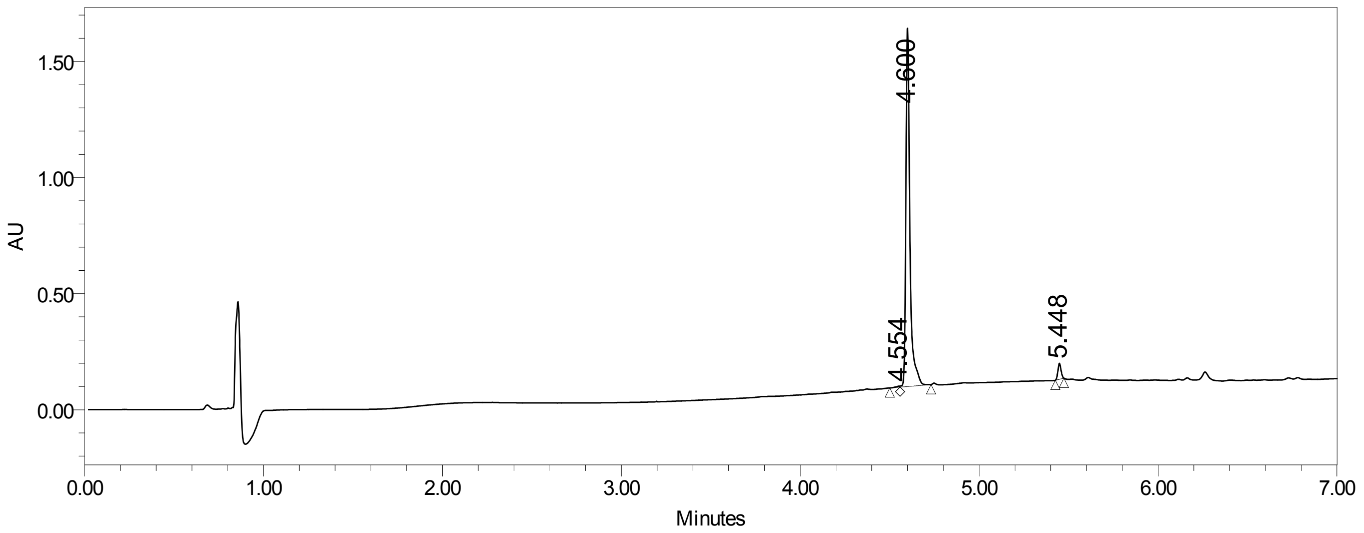

Supplementary Figure 4. α-VI UPLC purity analysis, UPLC gradient from 10-95% MeCN/$H_2O$ over 6 minutes (0.3 mL/min; column- Waters Acquity BEH C4 1.7 μm, 2.1 x 100 mm, purity >95%)

**α/β-VI-1**

Sequence: Ac-VA($\beta^3$L)DP($\beta^3$I)DIS($\beta^3$I)VL($\beta^3$N)KIK($\beta^3$S)DL($\beta^3$E)ESK($\beta^3$E)WI($\beta^3$R)RSN($\beta^3$Q)KL($\beta^3$D)SI-$NH_2$

Calculated monoisotopic [M+H]: 4346.5

Calculated monoisotopic [M+2H]: 2173.8

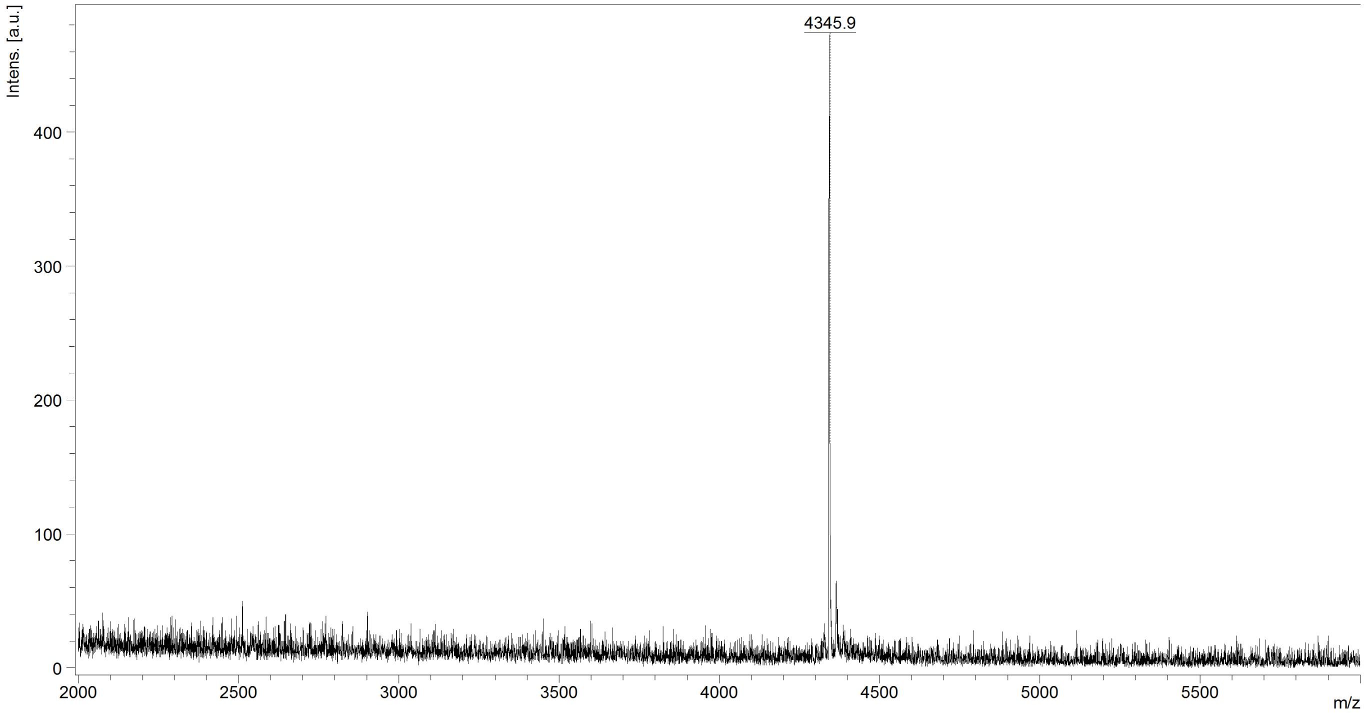

Supplementary Figure 5. α/β-VI-1 MALDI-TOF analysis

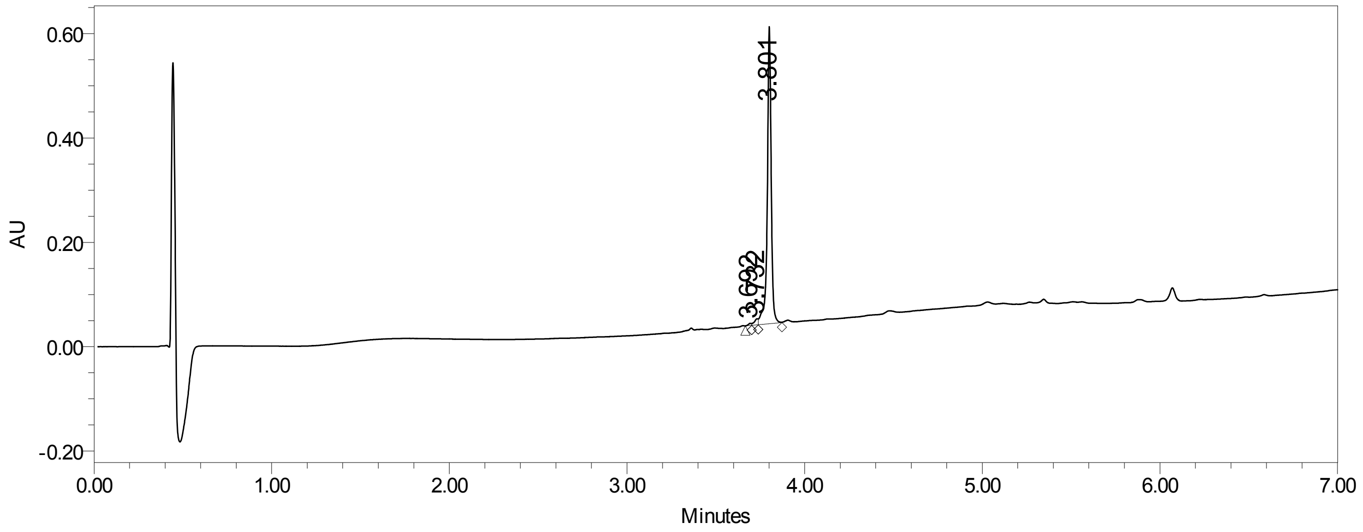

Supplementary Figure 6. α/β-VI-1 UPLC purity analysis, UPLC gradient from 10-95% MeCN/$H_2O$ over 6 minutes (0.3 mL/min; column-Waters Acquity BEH C4 1.7 μm, 2.1 x 100 mm, purity >95%)

**α/β-VI-2**

Sequence:

Ac-VA(ACPC)DP(ACPC)DIS(ACPC)VL(ACPC)KIK(ACPC)DL($\beta^3$E)ESK(APC)WI(APC)RSN(ACPC)KL($\beta^3$D)SI-$NH_2$

Calculated monoisotopic [M+H]: 4171.4

Calculated monoisotopic [M+2H]: 2086.2

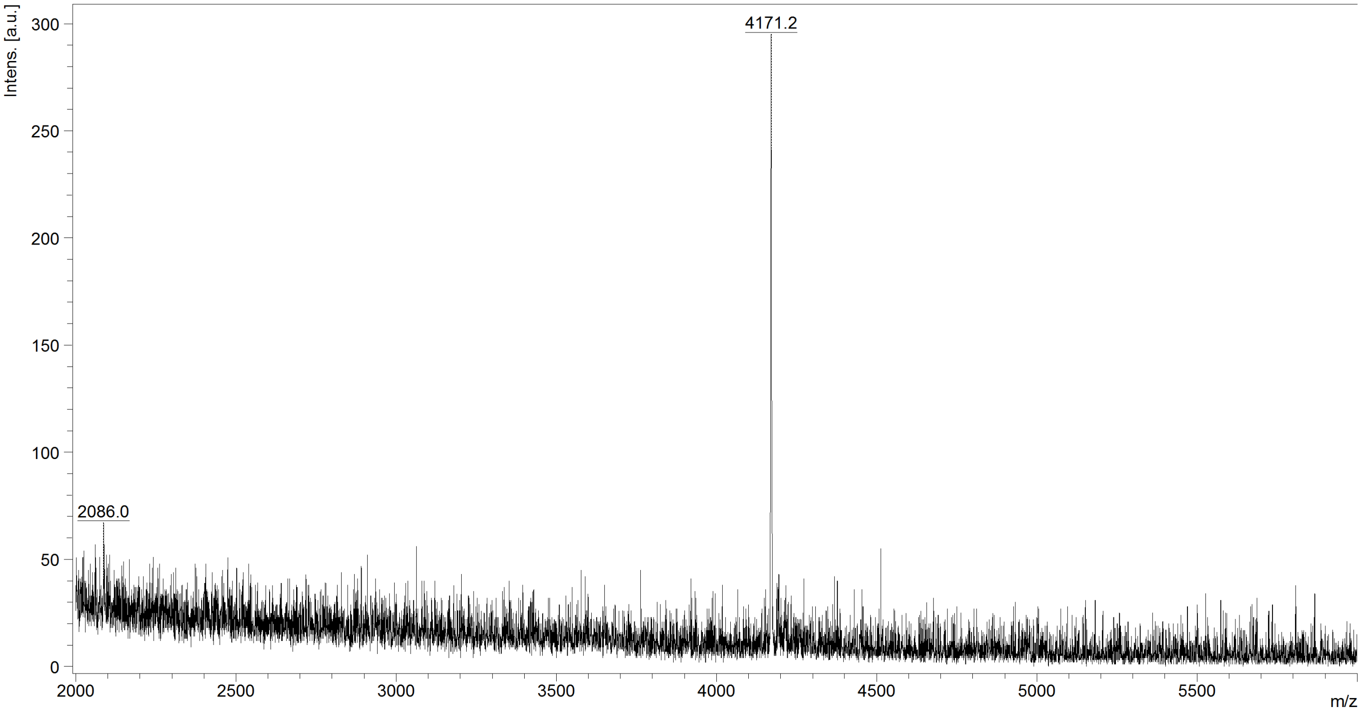

Supplementary Figure 7. α/β-VI-2 MALDI-TOF analysis

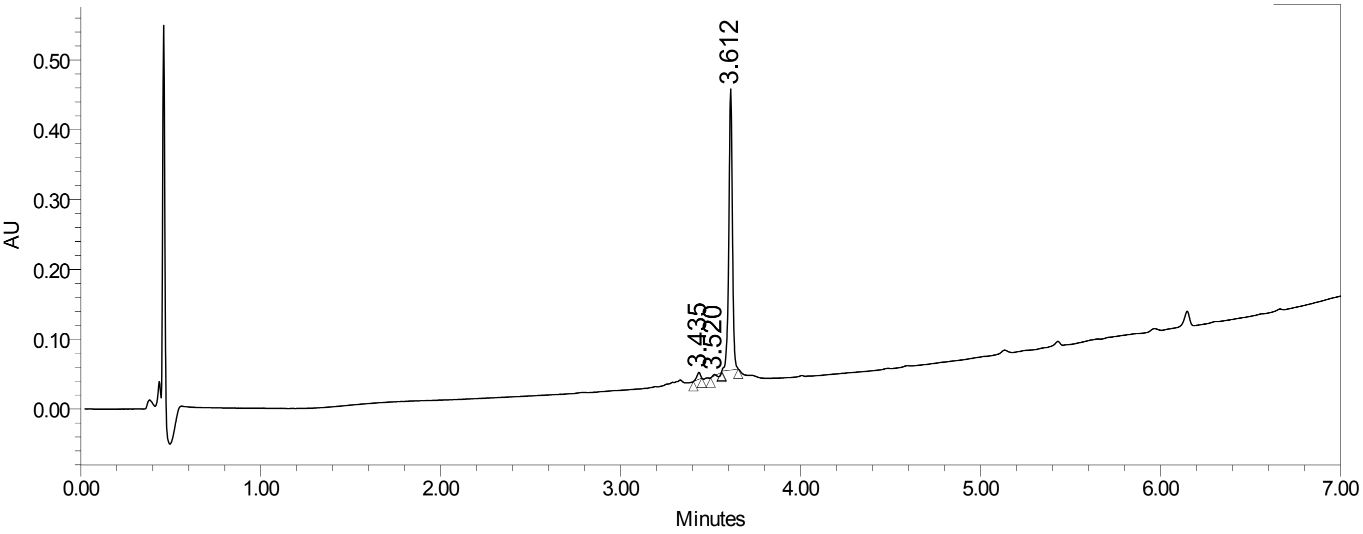

Supplementary Figure 8. α/β-VI-2 UPLC purity analysis, UPLC gradient from 10-95% MeCN/$H_2O$ over 6 minutes (0.3 mL/min; column-Waters Acquity BEH C4 1.7 μm, 2.1 x 100 mm, purity >95%)

**α/β-VI-3**

Sequence: Ac-VA($\beta^3$L)DPI($\beta^3$D)IS($\beta^3$I)VLN($\beta^3$K)IK($\beta^3$S)DLE($\beta^3$E)SK($\beta^3$E)WIR($\beta^3$R)SN($\beta^3$Q)KLD($\beta^3$S)I-$NH_2$

Calculated monoisotopic [M+H]: 4346.5

Calculated monoisotopic [M+2H]:2173.8

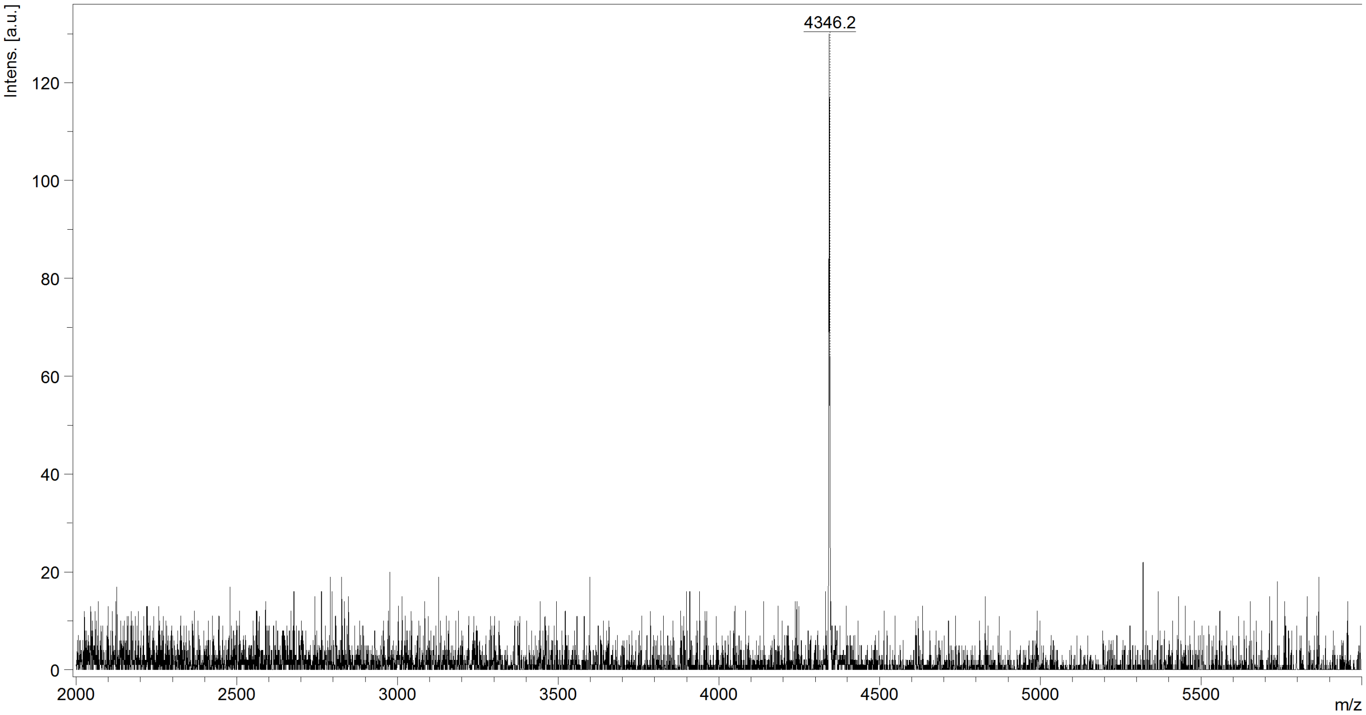

Supplementary Figure 9. α/β-VI-3 MALDI-TOF analysis

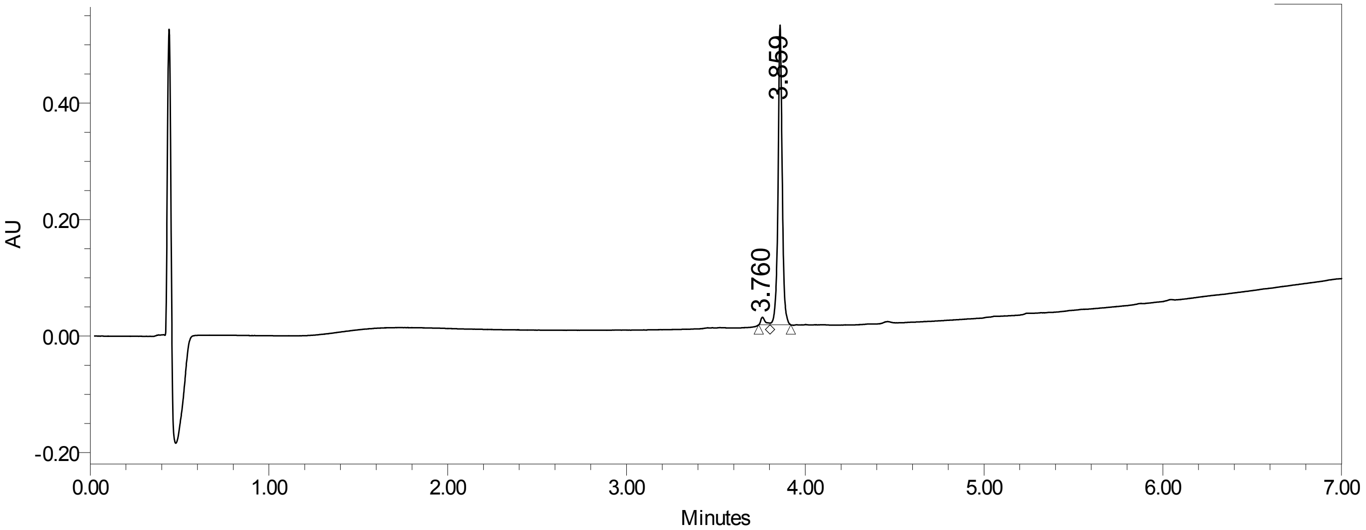

Supplementary Figure 10. α/β-VI-3 UPLC purity analysis, UPLC gradient from 10-95% MeCN/$H_2O$ over 6 minutes (0.3 mL/min; column-Waters Acquity BEH C4 1.7 μm, 2.1 x 100 mm, purity >95%)

**α/β-VI-4**

Sequence:

Ac-VA(ACPC)DPI($\beta^3$D)IS(ACPC)VLN(APC)IK(ACPC)DLE($\beta^3$E)SK($\beta^3$E)WIR(APC)SN(ACPC)KLD(ACPC)I-$NH_2$

Calculated monoisotopic [M+H]: 4185.3

Calculated monoisotopic [M+2H]: 2093.2

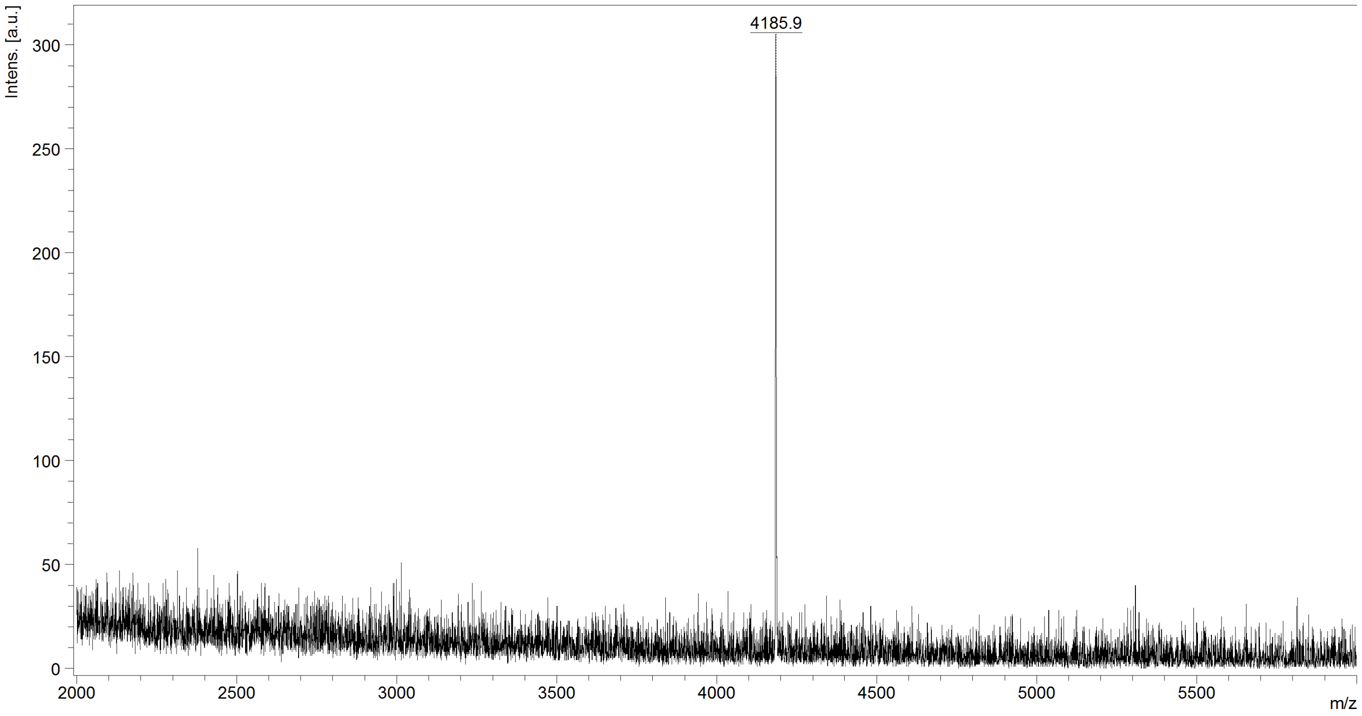

Supplementary Figure 11. α/β-VI-4 MALDI-TOF analysis

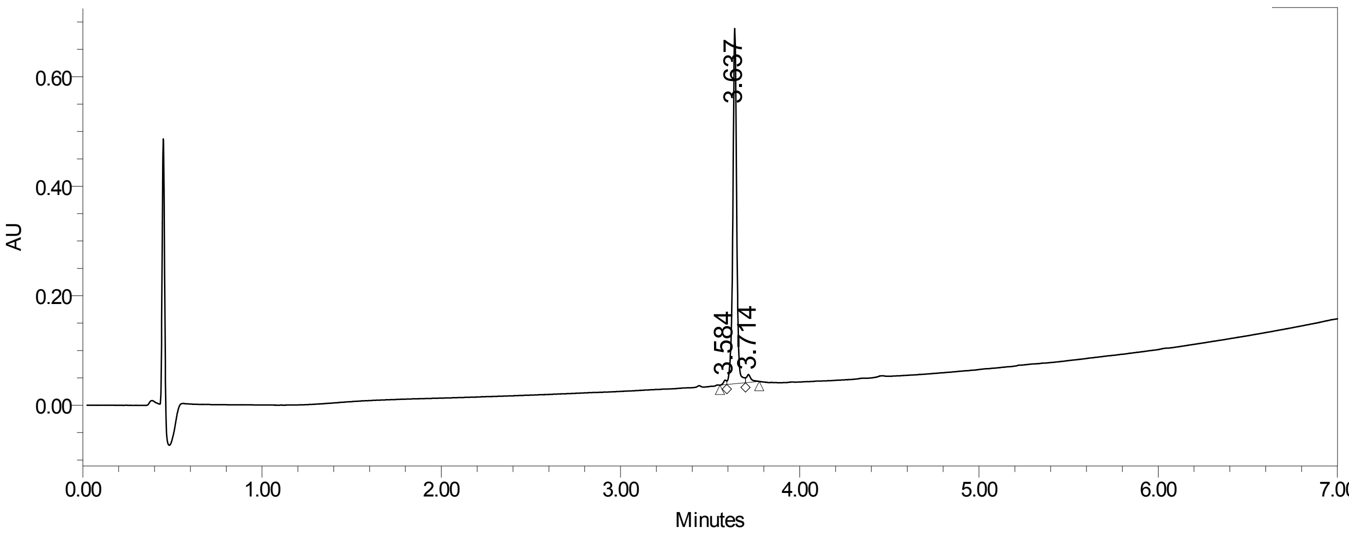

Supplementary Figure 12. α/β-VI-4 UPLC purity analysis, UPLC gradient from 10-95% MeCN/$H_2O$ over 6 minutes (0.3 mL/min; column- Waters Acquity BEH C4 1.7 μm, 2.1 x 100 mm, purity >95%)

**α/β-VI-5**

Sequence: Ac-VALDPIDIS(β3I)VL(β3N)KIK(β3S)DL(β3E)ESK(β3E)WI(β3R)RSN(β3Q)KL(β3D)SI-$NH_2$

Calculated monoisotopic [M+H]: 4318.4

Calculated monoisotopic [M+2H]: 2159.7

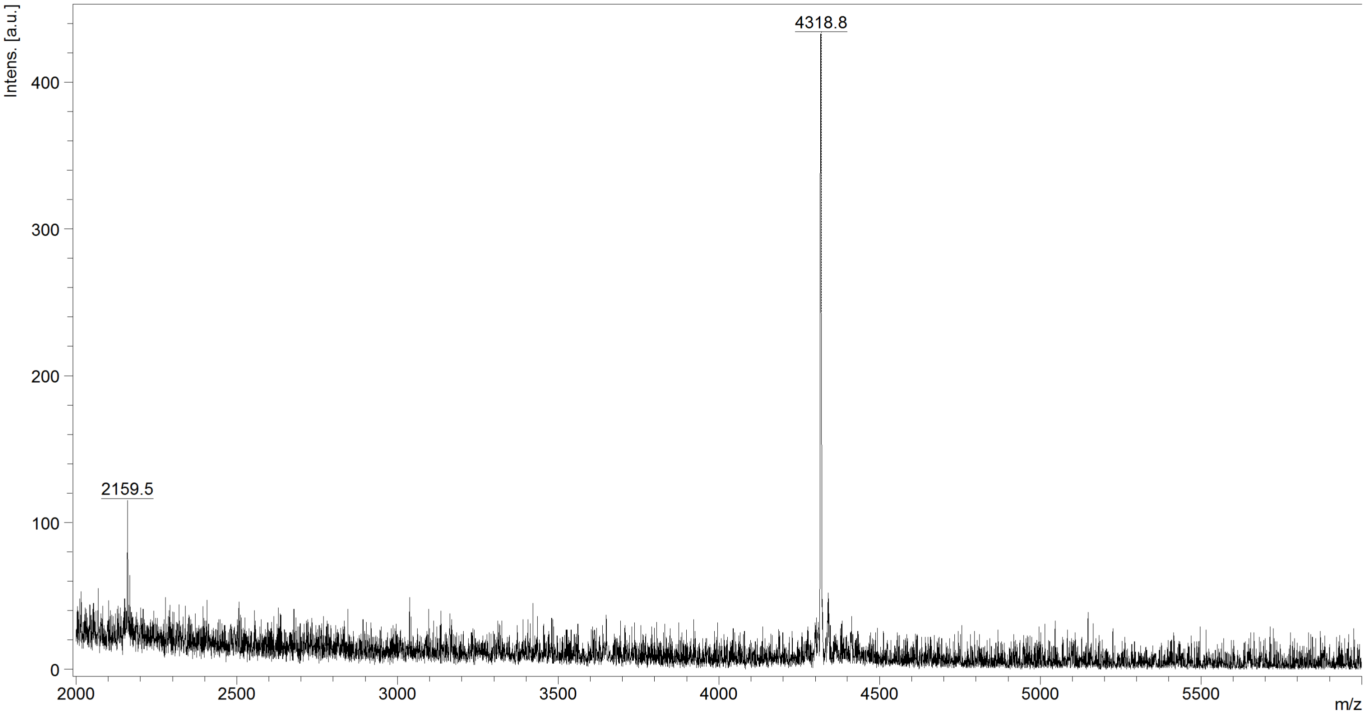

Supplementary Figure 13. α/β-VI-5 MALDI-TOF analysis

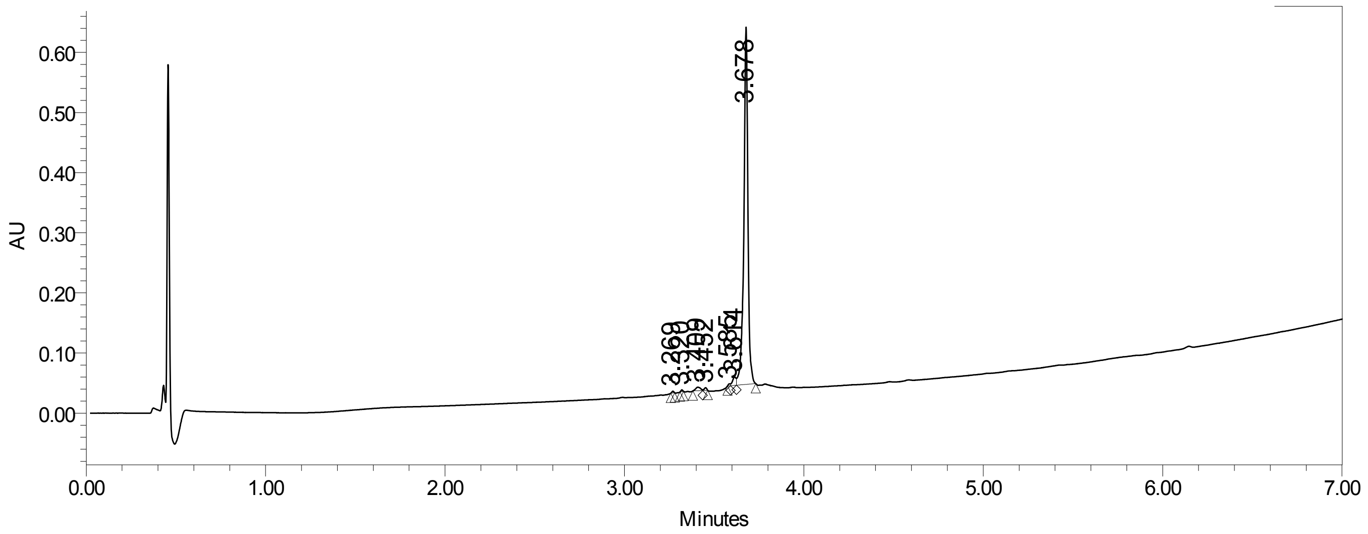

Supplementary Figure 14. α/β-VI-5 UPLC purity analysis, UPLC gradient from 10-95% MeCN/$H_2O$ over 6 minutes (0.3 mL/min; column-Waters Acquity BEH C4 1.7 μm, 2.1 x 100 mm, purity >95%)

**α/β-VI-6**

Sequence:

Ac-VALDPIDIS(ACPC)VL(ACPC)KIK(ACPC)DL($\beta^3$E)ESK(APC)WI(APC)RSN(ACPC)KL($\beta^3$D)SI-$NH_2$

Calculated monoisotopic [M+H]: 4175.4

Calculated monoisotopic [M+2H]: 2088.2

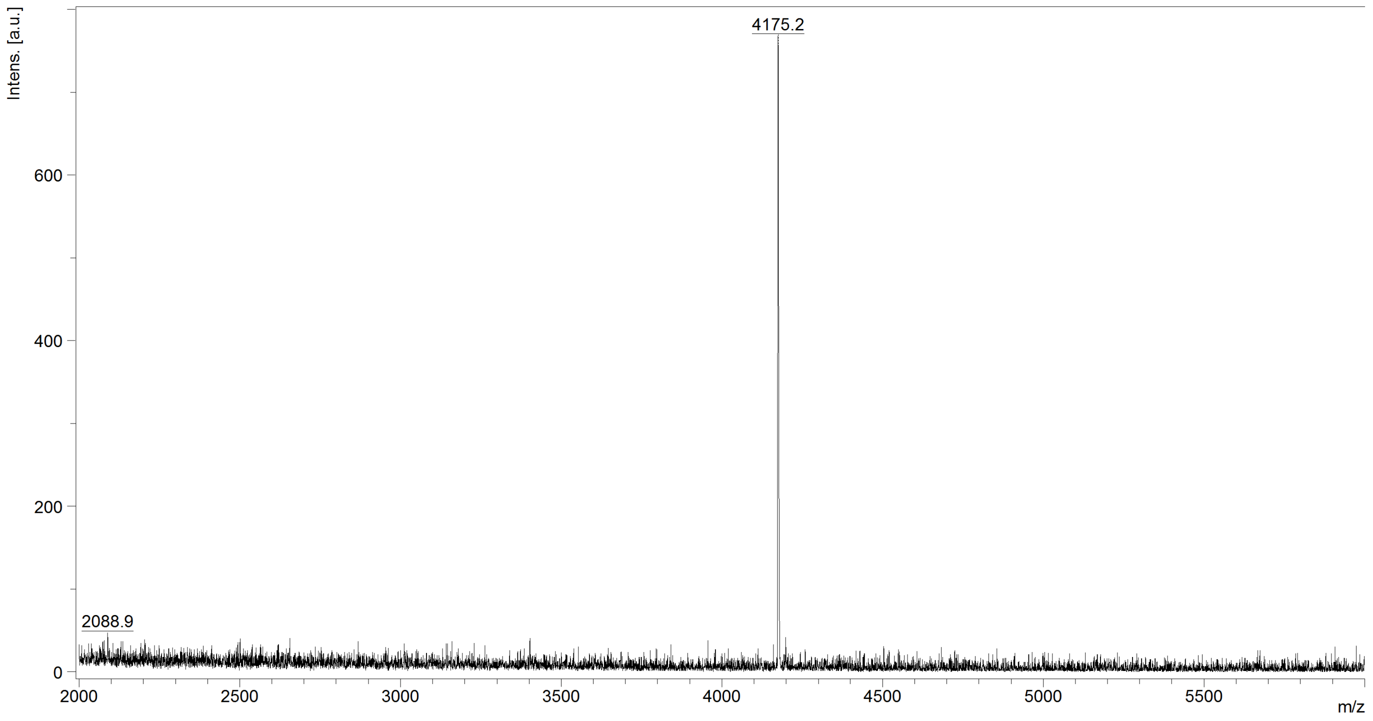

Supplementary Figure 15. α/β-VI-6 MALDI-TOF analysis

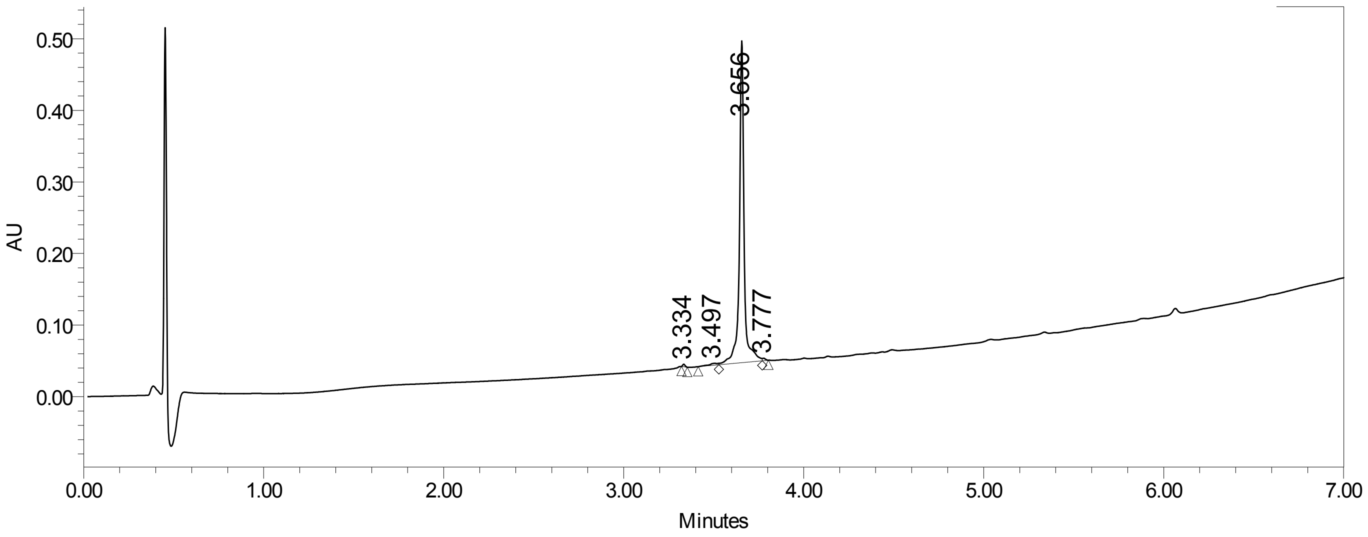

Supplementary Figure 16. α/β-VI-6 UPLC purity analysis, UPLC gradient from 10-95% MeCN/$H_2O$ over 6 minutes (0.3 mL/min; column- Waters Acquity BEH C4 1.7 μm, 2.1 x 100 mm, purity >95%)

**α/β-VI-7**

Sequence: Ac-VALDPIDIS($\beta^3$I)VLN($\beta^3$K)IK($\beta^3$S)DLE($\beta^3$E)SK($\beta^3$E)WIR($\beta^3$R)SN($\beta^3$Q)KLD($\beta^3$S)I-$NH_2$

Calculated monoisotopic [M+H]: 4318.4

Calculated monoisotopic [M+2H]: 2159.7

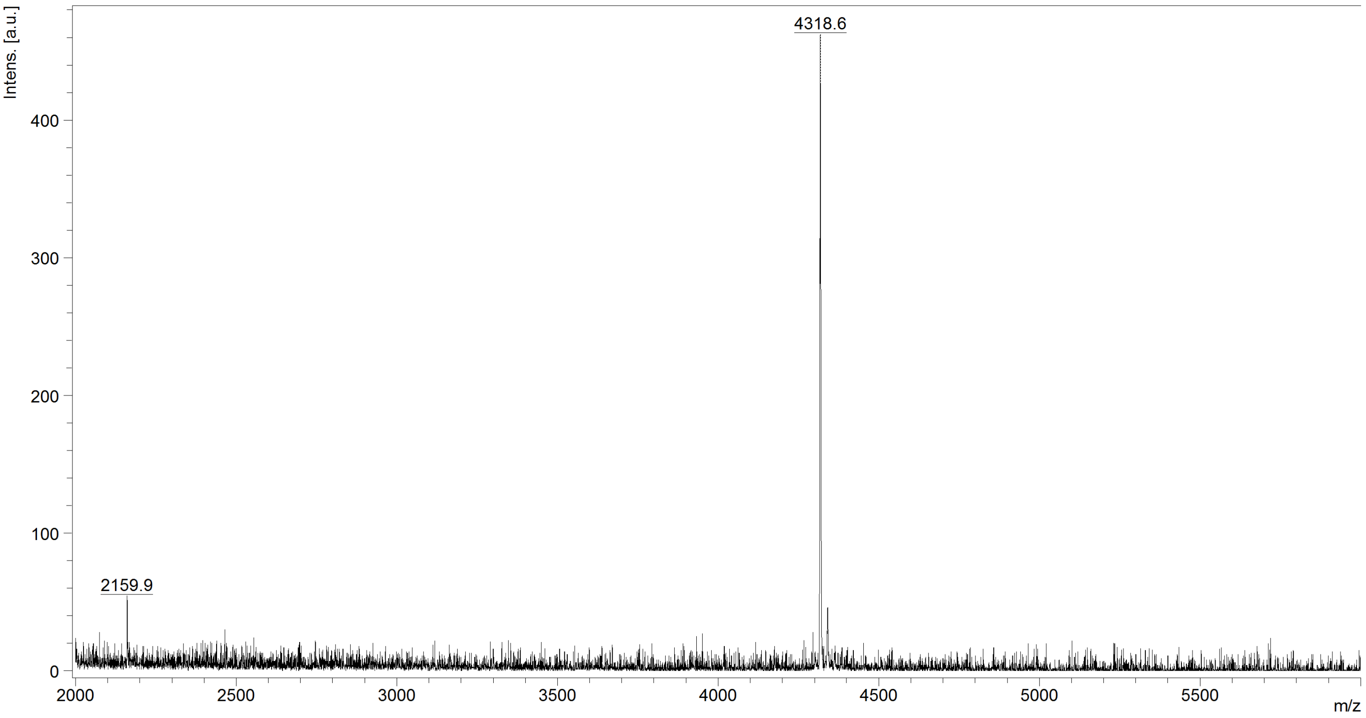

Supplementary Figure 17. α/β-VI-7 MALDI-TOF analysis

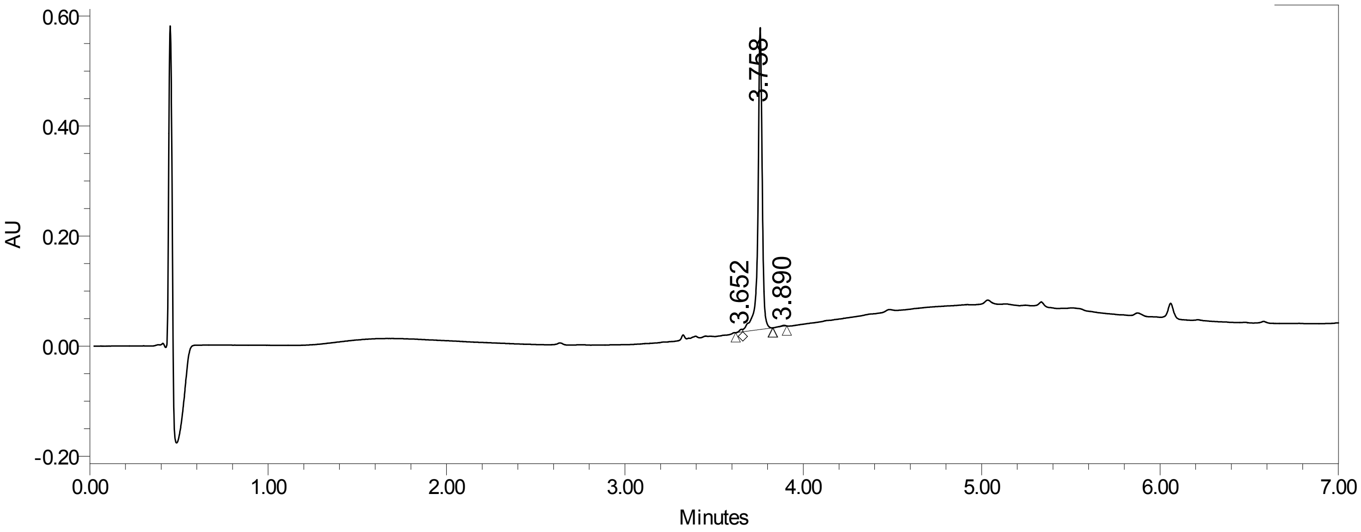

Supplementary Figure 18. α/β-VI-7 UPLC purity analysis, UPLC gradient from 10-95% MeCN/$H_2O$ over 6 minutes (0.3 mL/min; column- Waters Acquity BEH C4 1.7 μm, 2.1 x 100 mm, purity >95%)

**α/β-VI-8**

Sequence:

Ac-VALDPIDIS(ACPC)VLN(APC)IK(ACPC)DLE($\beta^3$E)SK($\beta^3$E)WIR(APC)SN(ACPC)KLD(ACPC)I-$NH_2$

Calculated monoisotopic [M+H]: 4203.3

Calculated monoisotopic [M+2H]: 2102.2

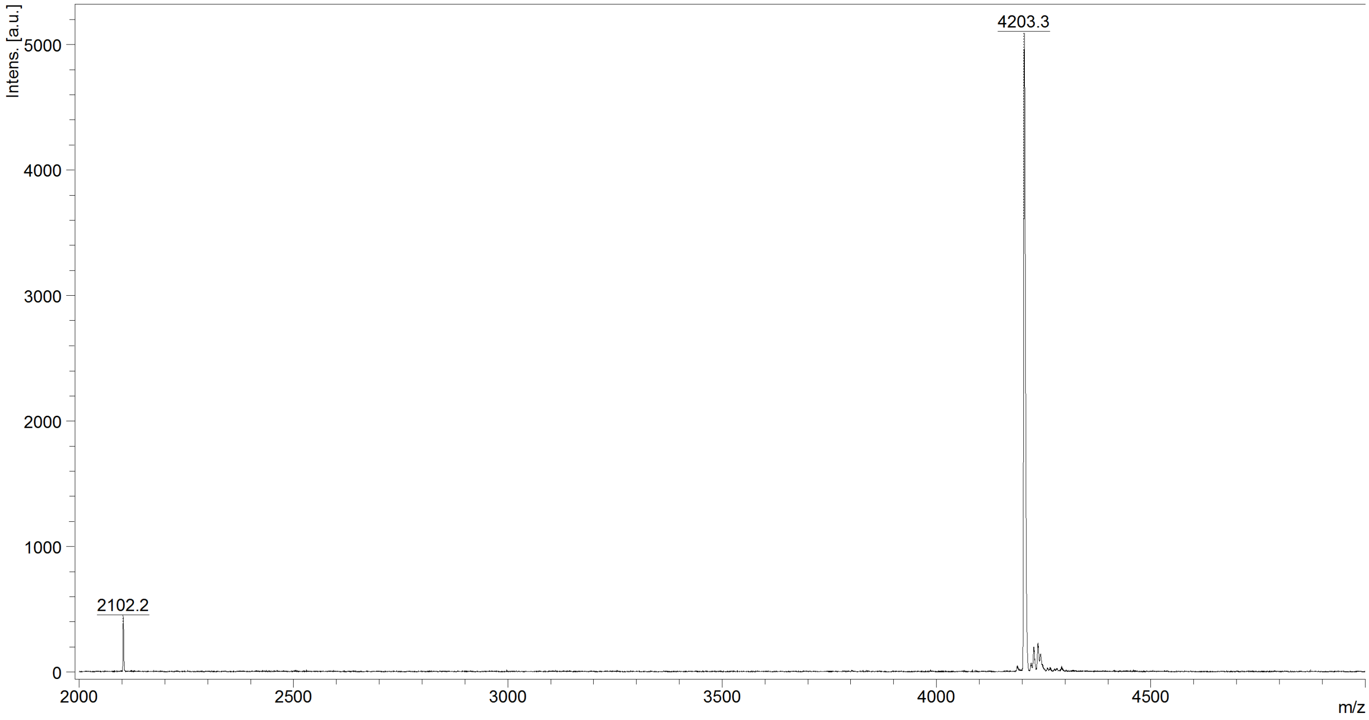

Supplementary Figure 19. α/β-VI-8 MALDI-TOF analysis

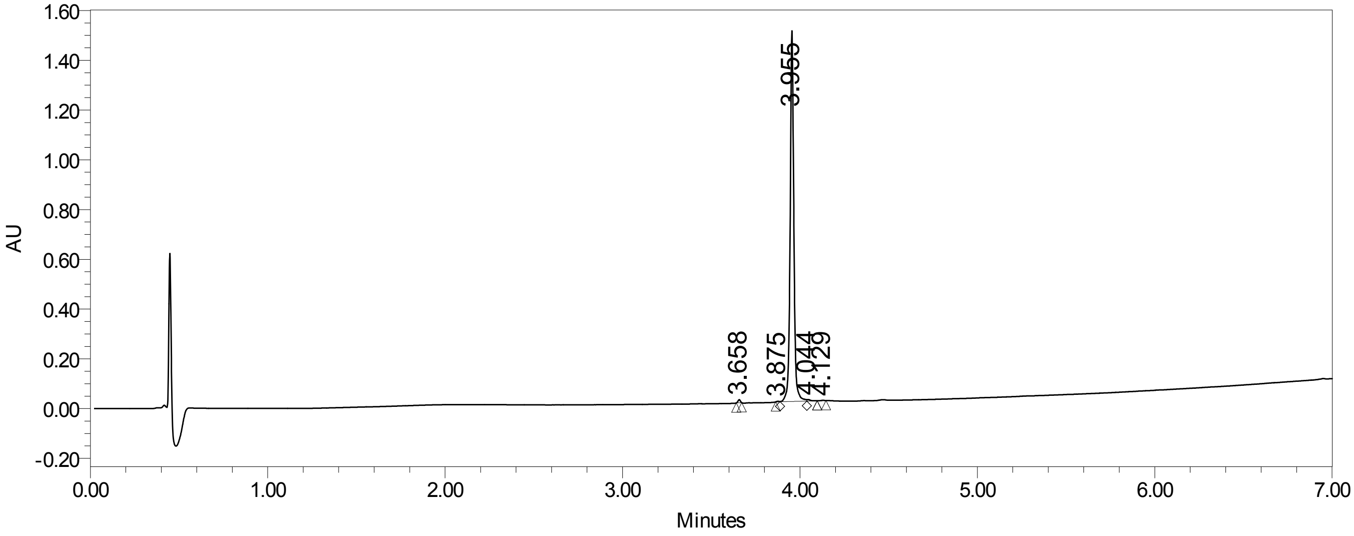

Supplementary Figure 20. α/β-VI-8 UPLC purity analysis, UPLC gradient from 10-95% MeCN/$H_2O$ over 6 minutes (0.3 mL/min; column-Waters Acquity BEH C4 1.7 μm, 2.1 x 100 mm, purity >95%)

### Circular Dichroism Spectroscopy

All circular dichroism (CD) spectroscopy experiments were performed on an Aviv Biomedical model 420 CD spectrometer. Samples were prepared in 1 mm quartz strain-free cuvettes (Hellma) using 50 μM peptide in phosphate buffered saline (PBS). Wavelength scans were collected from 260 nm to 200 nm with a 1 nm bandwidth and 10 second averaging time. The CD spectrometer was calibrated with a 1 mg/mL solution of camphor sulfonic acid (CSA) in water. For thermal denaturation experiments, ellipticity was measured at 222 nm as temperature was raised from 5–98 °C in 3-degree increments with a 5-minute equilibration time and a 10-second averaging time for each measurement.

#### Thermal Denaturation Assays for Specific HRN+HRC Pairs

In each thermal denaturation experiment, ellipticity of a 1:1 mixture of two peptides (50 μM each in PBS) was measured at 222 nm as temperature was raised from 5–98 °C in 3-degree increments with a 5-minute equilibration time and a 10-second averaging time for each measurement.

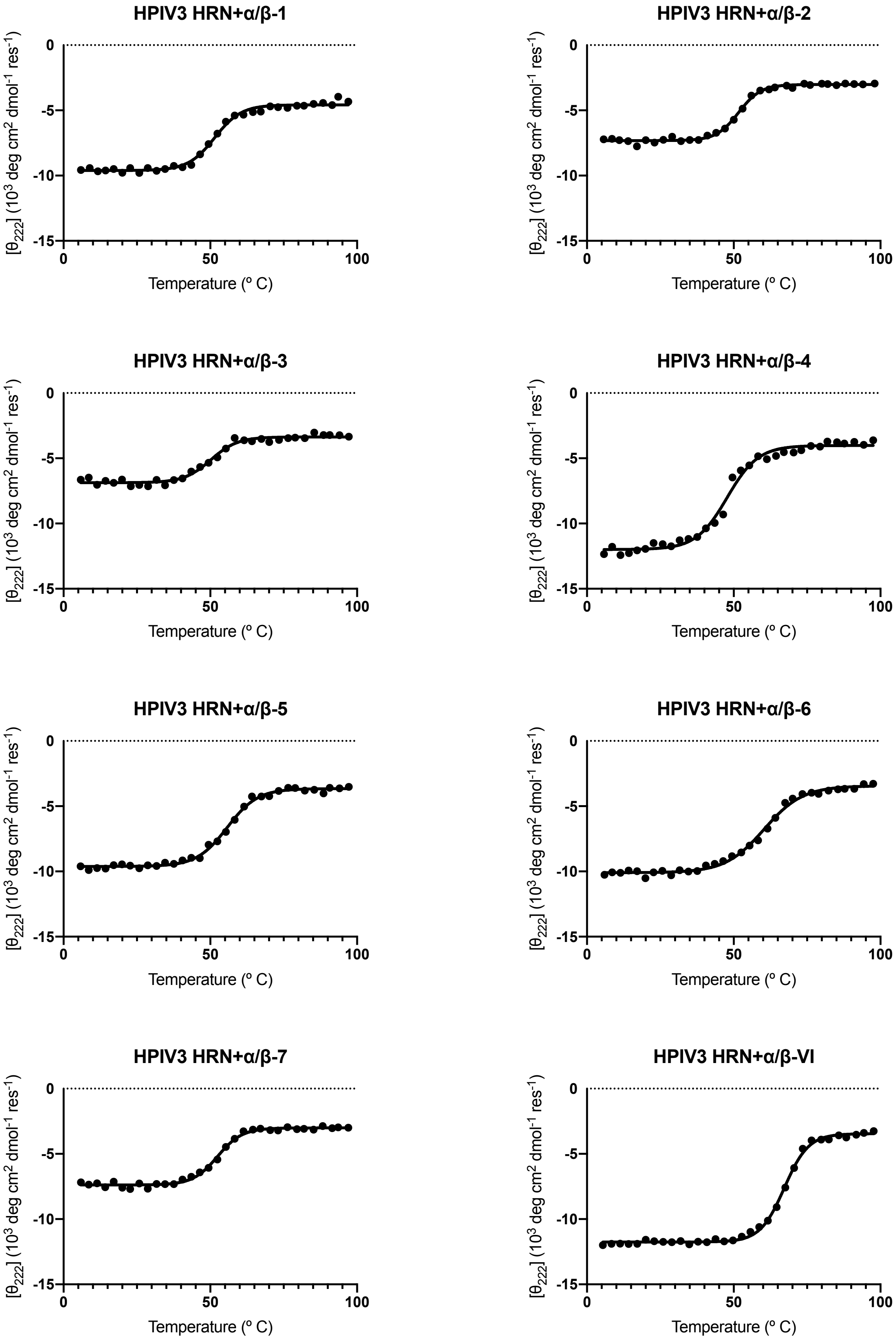

Supplementary Figure 21 Thermal denaturation spectra of co-assemblies between HPIV3 HRN and α/β-VI-1–8 peptides

### Synthesis of Peptide–Cholesterol Conjugates

#### Synthesis of α/β-VI-8–GSGSGC

A peptide corresponding to α/β-VI-8 with an additional C-terminal -GSGSGC sequence was synthesized by MA-SPPS.

Ac-VALDPIDIS(ACPC)VLN(APC)IK(ACPC)DLE($\beta^3$E)SK($\beta^3$E)WIR(APC)SN(ACPC)KLD(ACPC)IGSGSGC-$NH_2$

Calculated monoisotopic [M+H]: 4651.5

Calculated monoisotopic [M+2H]: 2326.3

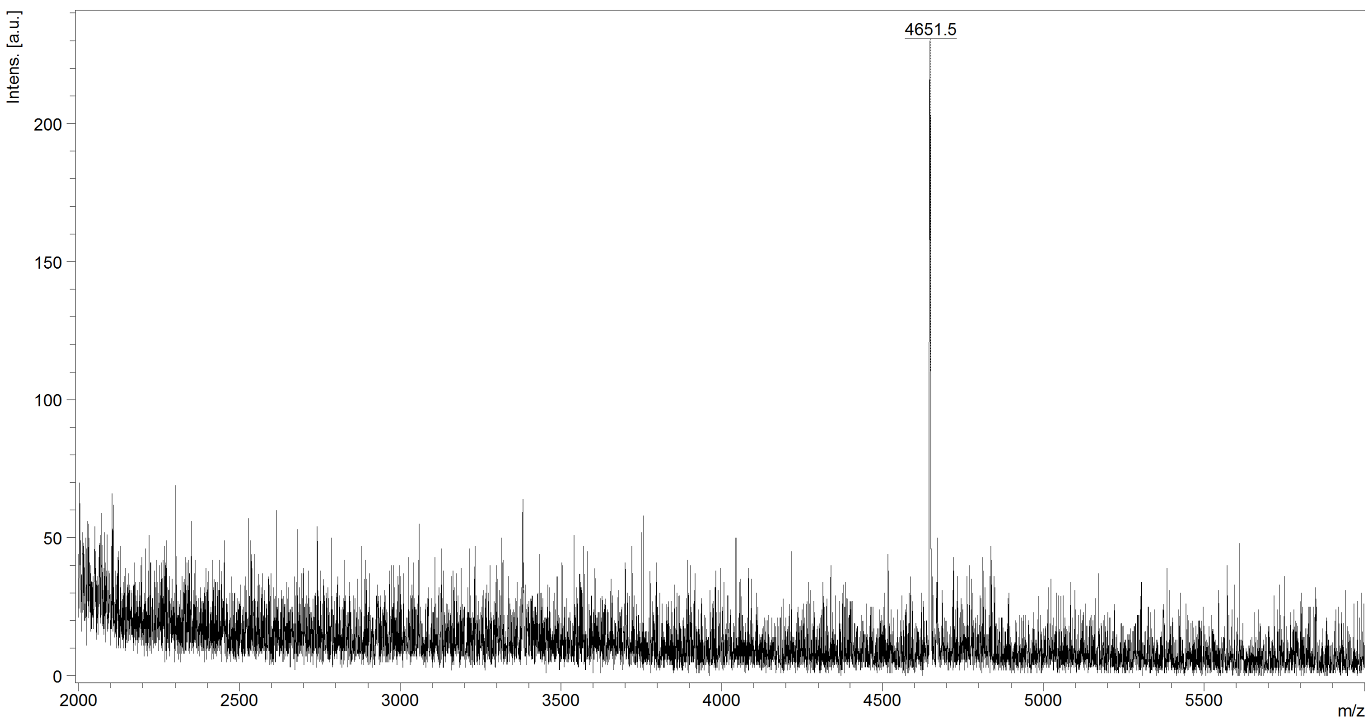

Supplementary Figure 22. α/β-VI-8-GSGSGC MALDI-TOF analysis

#### Synthesis of α/β-VI-8–$PEG_4$–Chol

The cholesterol conjugation was achieved using displacement of an α-bromoamide as shown in Fig S23. In a nitrogen-purged vial, BrAcNH-$PEG_4$-Chol reagent (43 mg, 57 µmol, 1.5 equiv.) was dissolved in 2.3 mL degassed DMSO. In a separate nitrogen-purged vial outfitted with a stir bar, Ac-α/β-VI-8-GSGSGC-$NH_2$ (178 mg, 38 µmol, 1.0 equiv.) was dissolved in 10.2 mL degassed DMSO. The BrAcNH-$PEG_4$-Chol solution was added to the peptide solution dropwise via syringe, followed by addition of 330 µL *N,N*-diisopropylethylamine. The reaction mixture was stirred for 90 minutes. Tris(2-carboxyethyl)phosphine (22 mg, 77 µmol, 2.0 equiv.) was added, and the mixture was stirred for an additional 15 minutes. The resulting peptide-cholesterol conjugate was purified by reverse-phase HPLC on a Shimadzu HPLC system (SCL-10VP system controller, LC-6AD pumps, SIL-10ADVP autosampler, SPD-10VP UV-vis detector, FRC-10A fraction collector) equipped with a Waters XSelect CSH Prep C18 column (5 µm particle size, 19 mm × 250 mm) using a gradient of 65-79% acetonitrile+0.1% trifluoroacetic acid in water+0.1% trifluoroacetic acid over 18 minutes at a flow rate of 12 mL/min.

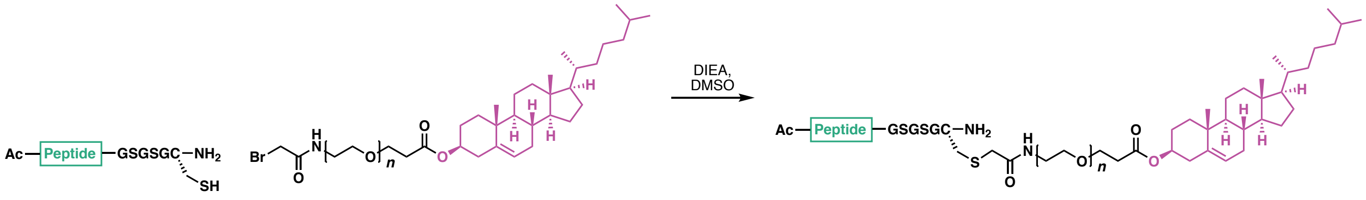

Supplementary Figure 23 Synthesis of cholesterol-conjugated peptides

Calculated monoisotopic [M+H]: 5325.0

Calculated monoisotopic [M+2H]: 2663.0

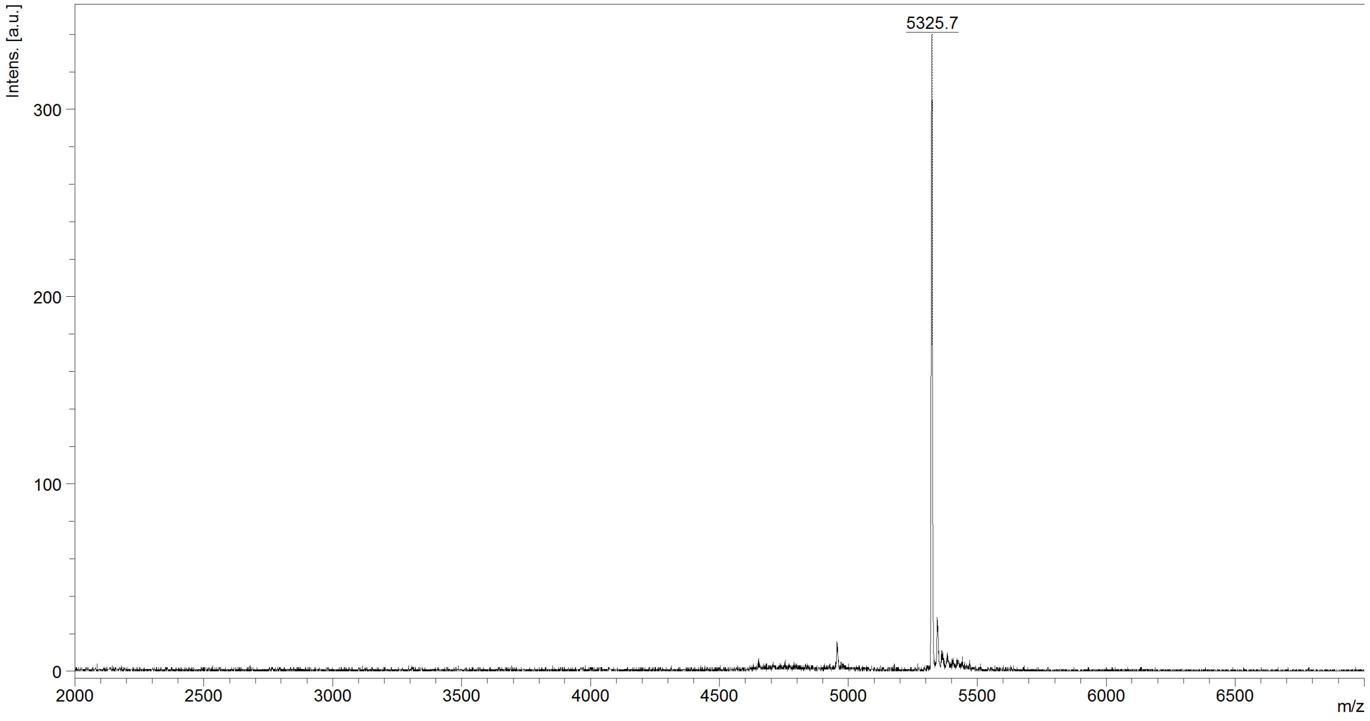

Supplementary Figure 24. α/β-VI-8–$PEG_4$–Chol MALDI-TOF analysis

### X-ray Crystallography

#### Crystallization Conditions

Separate stock solutions of the HPIV3 HRN and α/β-VI-8 peptides were prepared by dissolving the lyophilized peptide powder in water. Peptide concentration was measured by UV absorbance using tryptophan and tyrosine as the chromophores. Co-crystallization solutions were then prepared by mixing equal molar amounts of the relevant individual peptide solutions. Crystals were grown using hanging drop vapor diffusion. A 2 μL drop that comprised a 1:1 mixture of co-crystallization solution and the crystallization condition was placed on a glass cover slide that was then inverted to seal a well containing 150 μL of the crystallization condition. The crystallization conditions are given below.

- HPIV3 HRN+α/β-VI (α/β-8): 30 mM NaF, 30 mM NaBr, 30 mM NaI, 20% (v/v) PEG 500 MME, 10% (w/v) PEG 20000 in 100 mM imidazole/MES monohydrate buffer (pH 6.5)

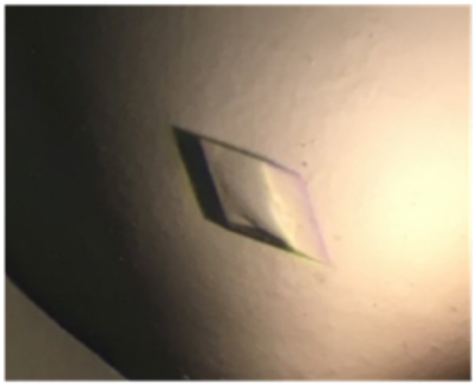

Supplementary Figure 25 Crystal morphology for HPIV3 HRN+α/β-VI-8

#### X-ray Data Collection

Crystals were looped and vitrified in liquid nitrogen. Diffraction data for HPIV3 HRN+α/β-VI-8 were collected using a Bruker AXS MICROSTAR generator housed and maintained at the University of Wisconsin in the lab of Professor Katrina Forest.

#### Data Processing, Structure Solution, and Refinement

Data were indexed and integrated using the program *XDS*. Data were then scaled and merged using the program *XSCALE*.[1] A molecular replacement solution was found with the program *Phaser* for each dataset using a poly-alanine dimeric model of the C-terminal and N-terminal heptad repeat regions from the postfusion HPIV3 F ectodomain (PDB: 1ZTM) as a search model. Model refinement was carried out using the program *phenix.refine* in combination with manual real-space model building and refinement in the program *Coot*.[2,3]

**Supplementary Table 2. HPIV3 HRN+ α/β-VI-8 (PDB: 6VJO)**

| | |
|---|---|
| **Data collection** | |
| X-ray source | Bruker AXS MICROSTAR |
| X-ray detector | Bruker SMART 6000 |
| Detector distance (mm) | 50 |
| Oscillation range (°) | 1.0 |
| Wavelength (Å) | 1.542 |
| Space group | R 3 |
| a / b / c (Å) | 48.3 / 48.5 / 134.5 |
| α / β / γ (°) | 90 / 90 / 120 |
| Matthews coefficient ($Å^3$/Da) | 3.07 |
| Solvent content (%) | 60.0 |
| Resolution range (Å) | 35.55–2.00 (2.13–2.00) |
| Number of observations | 66793 (3698) |
| Unique reflections | 7675 (1107) |
| Completeness | 96.8% (83.9%) |
| Redundancy | 8.7 (3.3) |
| Mean $I/\sigma(I)$ | 10.5 (2.4) |
| CC1/2* | 1.00 (0.43) |
| $R_{merge}$ | 0.161 (0.807) |
| $R_{meas}$ | 0.171 (0.944) |
| $R_{pim}$ | 0.058 (0.478) |
| Wilson B-factor ($Å^2$) | 16.48 |
| **Refinement statistics** | |
| Refinement program | Phenix.refine: 1.17.1_3660 |
| Resolution range (Å) | 35.55–2.00 (2.13–2.00) |
| No. of unique reflections used in refinement | 7660 |
| Completeness (%) | 96.7% (83.6%) |
| Reflections in cross-validation set | 776 |
| R-value (work) | 20.0 |
| R-value (free) | 22.6 |
| R-value (overall) | 20.2 |
| Mean ADP ($Å^2$) | 32.3 |
